# Small siphophage binding to an open state of the LptDE outer membrane lipopolysaccharide translocon

**DOI:** 10.1101/2025.07.07.663478

**Authors:** Emily Dunbar, Robert Clark, Arnaud Baslé, Shenaz Allyjaun, Hector Newman, Julia Hubbard, Syma Khalid, Bert van den Berg

## Abstract

Bacteriophages are bacterial viruses that provide alternatives to small-molecule drugs to combat infections by antibiotic-resistant bacteria. To infect a bacterial host, a phage needs to bind to the bacterial surface via receptor binding proteins (RBPs), which are critical for determining host specificity. For functionally important receptors, the RBP-receptor interaction could be exploited via phage steering, where emerging bacterial resistance due to receptor modification could make bacteria less fit or virulent. Despite this, relatively little is known about RBP-receptor interactions. Here we build on the recent discovery of coliphages that have the outer membrane (OM) lipopolysaccharide translocon LptDE as their terminal receptor and show via cryogenic electron microscopy (cryoEM) that, surprisingly, the RBP of the small siphophage Oekolampad binds to a hitherto unobserved, open state of LptDE. The open lateral gate of LptD is occupied by a β-strand peptide originating from the degraded N-terminal jellyroll domain of LptD, suggesting the possibility of LptD inhibition via peptidomimetics. A structure of LptDE in complex with the superinfection exclusion (SE) protein Rtp45 of the Oekolampad-related phage Rtp shows a mechanism of SE where Rtp45-induced conformational changes in LptD resulting from steric clashes preclude RBP binding. Finally, analysis of spontaneous Oekolampad-resistant *E. coli* mutants identifies mutations in LptD that abolish the LptDE-RBP interaction *in vitro*. SDS-EDTA sensitivity assays of the mutants show no major defects in LptDE function, suggesting that phage steering via LptDE might be challenging.

**Significance:** The outer membrane (OM) of Gram-negative bacteria is a protective barrier generated by lipopolysaccharide (LPS), a complex glycolipid that makes up the outer leaflet of the OM. Following its synthesis and transport from the inner membrane, LPS is inserted into the OM by LptDE, an OM protein complex essential for most Gram-negative bacteria. LPS insertion requires a hitherto unobserved, open state of LptDE. Here we report the unexpected finding that the receptor binding protein (RBP) of a bacteriophage binds to an open state of LptDE, allowing isolation and visualisation via cryogenic electron microscopy. The lateral gate of the LptDE-RBP complex is occupied by a β-strand peptide, suggesting that the LptDE of pathogenic bacteria could be inhibited by peptidomimetics.

## Introduction

The increased threat of bacterial multidrug resistance has resulted in a renewed interest in phage therapy to combat pathogenic bacteria, especially difficult-to-treat Gram-negatives such as *Pseudomonas aeruginosa* and *Acinetobacter baumannii* (1). For a bacteriophage to inject its genome and infect a cell, phage attachment (adsorption) to the host cell must take place. For most phages, this step requires phage receptor binding proteins (RBPs) located in the phage tail, which recognise and bind to specific bacterial surface molecules such as outer membrane proteins (OMPs), polysaccharides, and lipopolysaccharides (2). Initial reversible binding often precedes an irreversible binding step to a different, terminal receptor that results in phage DNA injection. Despite its importance for determining host specificity, relatively little is known about the molecular details of bacterial receptor recognition by phage RBPs, especially in the case of the abundant Siphophages, which have long and flexible, non-contractile tails and RBPs that often target OMPs. An exception is bacteriophage T5, which targets the *E. coli* FhuA ferrichrome OM TonB-dependent transporter (TBDT) as its terminal receptor via its RBP pb5, and which employs the small periplasmic lipoprotein Llp for superinfection exclusion (SE) (3–5)

As with conventional antibiotic treatment of bacterial infections, phage therapy inevitably leads to bacterial resistance. In the example of phage T5 above, resistance could arise via, for example, shutting down FhuA expression by *E. coli*. This would likely be tolerated well, given that *E. coli* has six TBDTs for the acquisition of different iron-siderophore complexes (6), and under most conditions, the loss of one or several of those transporters would not be expected to greatly affect fitness. This latter point is a key aspect of phage steering, a strategy where phage therapy relies on the generation of resistance that renders the surviving bacteria either less virulent, less fit, or both (7–9). In the case of FhuA and phage T5, such a scenario is unlikely but might occur if ferrichrome were the most abundant source of iron available to *E. coli* at an infection site. However, for obvious reasons, phage steering is likely to be the most efficient for phages that target essential OMPs. The expression of such proteins cannot be shut down, and the generation of resistance might result in impaired function of the essential OMP and lower fitness or virulence. Only two OMPs are known to be essential in all or most Gram-negative bacteria: the BamA component of the β-barrel assembly machinery (BAM) and the lipopolysaccharide translocon LptDE (10, 11). The latter is the destination of lipopolysaccharide (LPS) molecules that traverse from the inner membrane to the OM via the seven-membered Lpt (lipopolysaccharide transport) system (12). This consists of the LptBCFG ABC transporter that extracts the LPS molecule from the IM and hands it off to the periplasmic LptA protein (13, 14). Several copies of LptA link LptBCFG to LptDE in the OM, forming a protein bridge spanning the periplasmic space that functions as an ATP hydrolysis-driven conveyor belt for LPS molecules (15, 16). Once they arrive at the LptDE complex, the LPS molecules are inserted into the outer leaflet of the OM (17, 18) via a process for which the details are not clear, but which must involve a separation of the first (S1) and last (S26) β-strands of the LptD barrel at the front of the complex to allow the translocating LPS molecule to move laterally into the OM from the lumen of the LptDE complex (19, 20). So far, in contrast to BamA, there is no structural information available for an open state of LptDE. Moreover, phages targeting LptDE were only identified recently, when a library of 68 novel coliphages (named the BASEL collection) was characterised together with several model phages and phage OMP receptors were identified via a panel of single OMP deletion strains (21). This approach failed for several small siphophages, but their receptor was subsequently identified as LptD via whole-genome sequencing of mutant strains that developed spontaneous resistance to those phages (21). The discovery of LptD-targeting phages provides opportunities to investigate whether an essential OMP could potentially be exploited towards phage steering.

Here we report the generation and cryo-EM structure determination of a complex between *Shigella flexneri* LptDE (SfLptDE; > 99% sequence identity with *E. coli* LptDE) and the RBP of the BASEL collection phage Oekolampad (bas018), which has LptD as terminal receptor (21). Oekolampad RBP (RBP_oeko_) binds to the extracellular face of an LptDE complex in which the LptD barrel has dramatically expanded at the front to form a wide channel connecting the periplasmic space to the extracellular environment. Moreover, the LptD lateral gate is occupied by clear density for a 12-residue β-strand peptide that originates from the N-terminal jellyroll domain of LptD, suggesting the potential for LptDE inhibition via peptidomimetics. The structure suggests that Oekolampad and similar small siphophages bind to an active, open state of the LptDE complex. To investigate the SE mechanism, we also determined the cryo-EM structure of the complex of SfLptDE with the SE lipoprotein Rtp45 of phage Rtp (22), which is related to Oekolampad. This structure shows that Rtp45 binds to closed LptDE at the periplasmic OM interface and causes conformational changes in LptD extracellular loops (EL) 4 and 5 that would abolish RBP binding. RBP_oeko_ binding to SfLptDE and subsequent infection is abolished by several point mutants of LptD. Interestingly, several of these mutants cause minimal changes in LptD and do not appear to increase the sensitivity of *E. coli* for OM stress, suggesting that LptDE functionality is not impaired and that phage steering via targeting LptDE might be challenging.

## Results

### The cryo-EM structure of SfLptDE contains bound LptM

We obtained a complex of *Sf*LptE with *E. coli* LptD following expression of the *imp4213* variant (23) of *Sf*LptDE (LptD lacking residues 330-352 in extracellular loop 4; see Methods, Figs. S1 and S2 and Table S1). The LptDE complexes from *Shigella* and *E. coli* are virtually identical in sequence (3 different residues in LptD and 1 different residue in LptE), and we consider this hybrid structure the same as *Sf*LptDE. We were able to obtain good-quality cryoEM maps around 3 Å resolution (Fig. 1A and Fig. S1) without having to resort to fiducial markers such as the Pro-macrobodies used for the recent determination of the cryo-EM structure of LptDE from *Neisseria gonorrhoeae* (NgLptDE) (24). Buildable density is visible for the entire LptDE complex, except for residues Tyr38-Ala67 of LptD and the periplasmic C-terminus of LptE (Ser169-Asn193) (Fig. 1A, B). Strikingly, additional density is visible close to periplasmic LptD loops and within the lumen of the complex (Movie S1). Based on the literature as well as on proteomic analysis of the sample, we assigned this density to the small lipoprotein LptM (formerly YifL; Uniprot P0ADN6). This protein was recently identified as a disulphide bond maturation factor of LptD and proposed to mimic bound LPS substrate (25, 26). Density is lacking for the Ala31-Thr44 segment of LptM, likely due to it being in the periplasmic space (Fig. 1C), but the assignment of the rest of the protein is unambiguous. Parts of all three acyl chains of the N-terminal lipid anchor are visible, protruding into the DDM micelle via the LptD intramembrane hole previously identified (27), supporting the notion that LptM mimics LPS. The LptM region between Gln45 and Asn54 interacts extensively with LptE, analogous to the proposed interaction of LptE with translocating LPS (Movie S1) (28). Following this region, the LptM chain is sandwiched between the functionally important LptD extracellular loop L4 (EL4) and the LptD C-terminus, before doubling back towards its own N-terminus, with intramolecular backbone hydrogen bonds between Lys23 and Ser63 (Fig. 1C). Compared to the AF2 prediction of bound LptM by Yang et al. (25), the positions of Cys20-Pro30 are very similar to the experimental structure, but beyond Gln45 both structures differ substantially, as evidenced by a difference of ∼20 Å between the C-terminal residue Tyr67 (Fig. 1D). This is consistent with lower confidence values of the AF2 prediction for the C-terminal part of LptM (25). Of note, no populations of LptDE complexes lacking LptM were observed.

**Figure 1.**
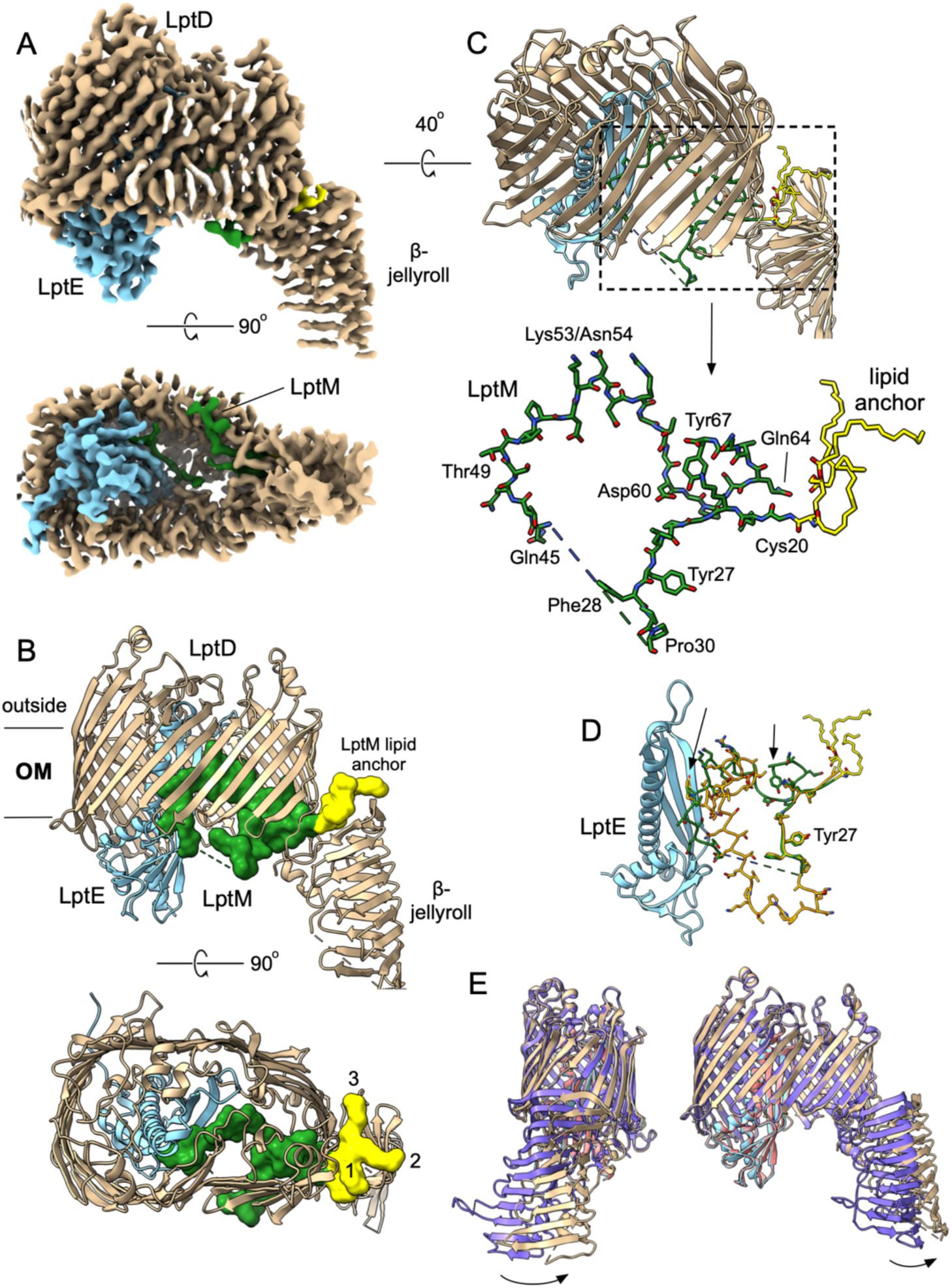
Cryo-EM structure of the LptDE-LptM complex. (A) Electron density maps shown at relatively high contour levels, with LptD coloured tan, LptE light blue and LptM dark green. The N-terminal lipid anchor and Cys20 of LptM are yellow. (B) Corresponding cartoon models coloured as in (A) and with LptM in surface representation. In the bottom panel, the acyl chains of the lipid anchor are numbered. (C) Stick model of LptM shown within the LptDE complex and without (bottom panel). The dashed line represents the Ala31-Thr44 segment, which lacks clear density. (D) Comparison of LptM with the predicted and minimised AF2 model in orange (Yang 2023). For reference, LptE is shown as well. Structures were superposed on LptD and apart from LptM, the structural differences are negligible. The arrows point to the C-terminal Tyr67 LptM residues, highlighting the large differences between the experimental LptM structure and the predicted model. (E) Superposition (on LptD) of the cryo-EM structure (coloured as above) with the X-ray crystal structure of SfLptDE, with LptD coloured purple and LptE pink (PDB 4Q35; Dong 2014). The arrows highlight the different positions of the N-terminal jellyroll domains. LptM, present only in the cryo-EM structure, is not shown for clarity.

Compared to the X-ray crystal structure (PDB 4Q35), the LptDE cryo-EM structure is largely identical except for the N-terminal jellyroll domain (Fig. 1E). In the cryo-EM structure, the jellyroll domain has moved/rotated approximately as a rigid body, breaking interactions between the barrel and the jellyroll domain (25). The difference may be caused by LptM, which is absent from the crystal structure. Indeed, the intramembrane hole in the crystal structure is too small to accommodate the LptM acyl chains. Moreover, in the crystal structure, the different LptDE monomers pack via end-to-end stacking of the jellyroll domains, and it is possible that this has induced a different jellyroll domain position in the crystal. Given our data suggest LptDE purifies with tightly bound LptM, this suggests that the crystallisation process may have selected for apo-LptDE. Interestingly, however, a crystal structure for *Pseudomonas aeruginosa* LptDE (expressed in *E. coli*) was recently published (PDB ID 8H1R; (26)) in which density for *E. coli* LptM residues Cys20-Asp32 is visible at the same location as in our cryo-EM structure. In addition, the recent cryo-EM maps of NgLptDE (24) also contains density consistent with bound LptM, although it was not modelled.

### Oekolampad RBP binds to an open state of the LptDE complex

Based on Maffei *et al*., we initially focused on the RBP of the coliphage Rtp, named Rtp44, which was proposed to be the terminal receptor for LptD based on its genomic location and homology (96% sequence ID) to the LptD-targeting AugustePiccard (bas01) RBP (21). However, *E. coli* expression of this protein failed. Switching targets, we obtained high expression levels for C-terminally His6-tagged RBP from Oekolampad (bas018; genus *Dhillonvirus*), which receptor was also identified as LptD by Maffei et al. RBP_Oeko_ has 33% sequence identity to Rtp44 and a very similar predicted fold (RMSD 1.1 Å for 244 Cα pairs out of 309 total; Fig. S3). This structural similarity is consistent with broader conservation among LptD-targeting RBPs, AlphaFold2 predicted models of RBPs from diverse LptD-dependent phages, including the putative RBP from OzMSK, reveal significant structural homology despite low sequence identity (29). Incubation of dodecyl-maltoside (DDM)-purified SfLptDE with ∼3-fold molar excess of RBP_Oeko_ resulted in the slow formation of a complex as judged by size-exclusion chromatography (SEC) in DDM (Fig. 2A). Given that complex formation is not quantitative, a larger-scale 48-hour co-incubation was set up using tag-less SfLptDE (Materials and Methods) and His-tagged RBP_Oeko_. The complex was purified via IMAC and SEC (Fig. 2B), and cryo-EM data were collected. Data processing gave two major particle classes (Table S1), the largest of which corresponded to a monomer and the other to a side-by-side dimer (Fig. S4). No density was observed for the LptD jellyroll domain in both classes, despite most of LptD being intact (Fig. 2A).

**Figure 2.**
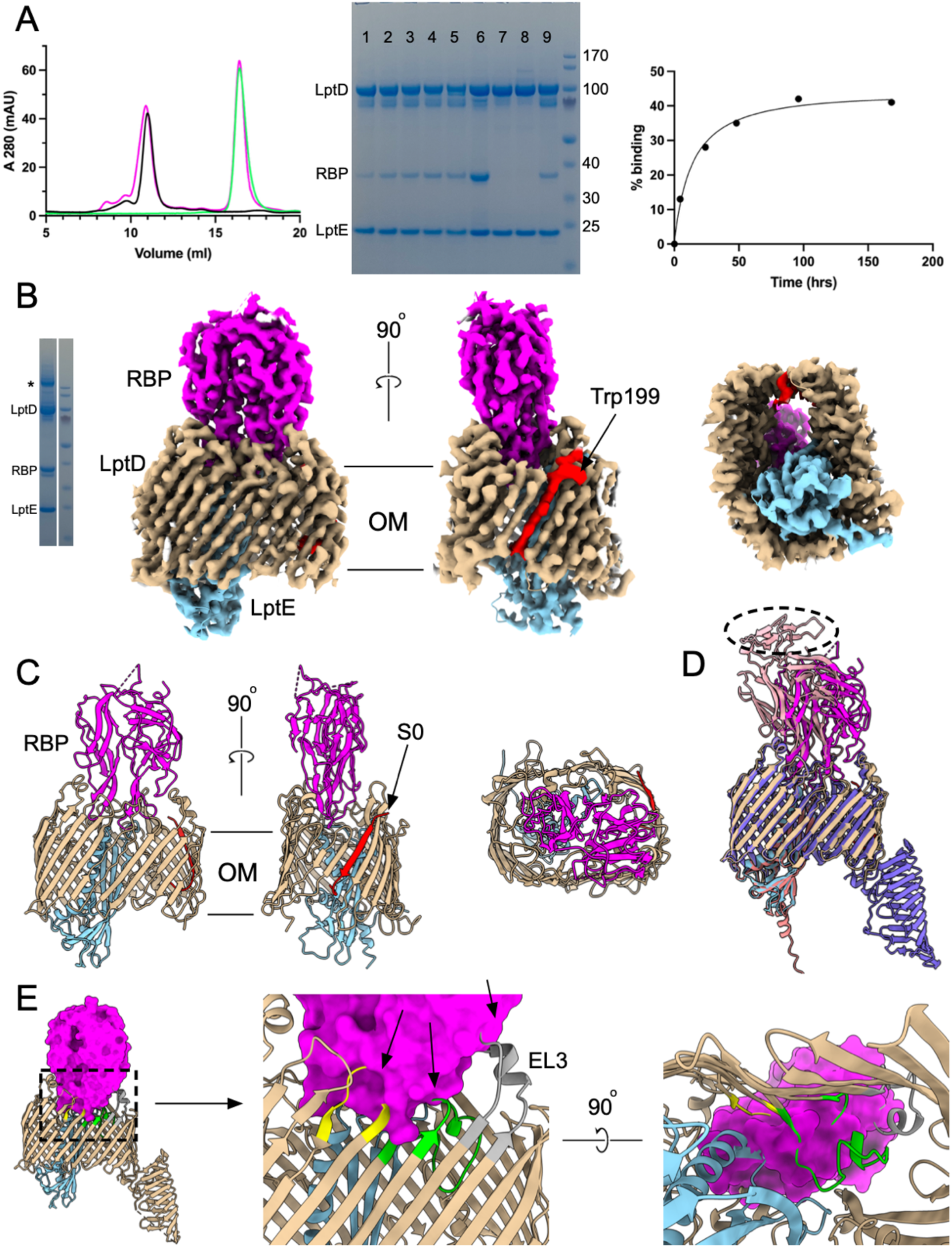
Generation and structure determination of the SfLptDE-RBP_Oeko_ complex. (A) Left panel, Representative SEC profiles in DDM for SfLptDE (black), RBP_Oeko_ (green) and the SfLptDE-RBP_Oeko_ complex (magenta; 3-fold molar excess of RBP). Samples were analysed after 48 hrs incubation at 4°C. Middle panel, SDS-PAGE samples of a time course of SfLptDE-RBP_Oeko_ complex formation. Lanes 1-5, incubations for 4, 24, 48, 96 and 168 h; lane 7, 1:1 molar ratio of SfLptDE:RBP_Oeko_; lane 8/9, SfLptDE Y671N mutant (R2) incubated with RBP_Oeko_ for 48 h in the presence (8) and absence (9) of TCEP; 10, SfLptDE wild type incubated with RBP_Oeko_ for 48 h. To enrich for the complex, the first half of the SfLptDE peak was collected and analysed (note the slight shift in Fig. 2A). Right panel, percentage RBP_Oeko_ binding generated from the SDS-PAGE samples via band densitometry. LptE was used as the reference band. The amount of complex formation after 48 h varied between 15-35%, depending on the LptDE preparation. (B) SDS-PAGE and cryo-EM density maps of the SfLptDE-RBP_Oeko_ complex at high contour. The * in the gel indicates disulphide-bonded, dimeric LptD. The rightmost panel shows a view from the periplasmic side. RBP is coloured magenta. The bound lateral gate peptide at the front of the complex is red, with the arrow indicating the putative Trp residue. (C) Cartoon models of SfLptDE-RBP_Oeko_, with the rightmost panel an extracellular view. (D) Comparison between the experimental and AF3-predicted SfLptDE-RBP_Oeko_ structures. RBP_Oeko_ of the AF3 model is coloured pink. The hatched oval represents the part of RBP_Oeko_ that is absent or very weak in the cryo-EM density. (E) Superposition (on LptD) of the RBP_Oeko_-LptDE complex (LptDE not shown for clarity) with the published LptDE crystal structure, showing extensive clashes of RBP with LptDE. The arrows point to parts of LptD EL3-5 that would overlap with bound RBP. EL3 is grey, EL4 lime green and EL5 yellow.

The monomeric particle class gave maps to ∼2.8 Å resolution and showed clear density for RBP_Oeko_ bound to the extracellular face of LptDE (Fig. 2B). RBP_Oeko_ inserts deeply into the LptDE lumen with a total interface area of almost 1700 Å^2^ as analysed via PISA (30), with a complex formation significance score (CSS) of 1.00. RBP_Oeko_ forms 20 hydrogen bonds and 2 salt bridges with LptD and 2 hydrogen bonds with LptE, one of which is between main-chain atoms. Thus, despite the very slow rate of complex formation, once established the interaction between LptDE and RBP_Oeko_ is likely to be tight. Most polar interactions of RBP_Oeko_ are with LptD EL3-5 and EL11. The AF3 prediction of RBP_Oeko_ in isolation is virtually identical to that in the complex structure (Cα RMSD 0.75 Å), suggesting that binding to LptDE causes very limited structural changes in RBP_Oeko_. Despite this, the prediction of the SfLptDE-RBP_Oeko_ complex shows a substantially different interaction, with RBP_Oeko_ shifted and inserted less deeply (Fig. 2D). In addition, the part of RBP_Oeko_ that would be proximal to the rest of the phage tail is very poorly ordered, an intriguing similarity to the phage T5 RBP (pb5) bound to the FhuA receptor (3). A superposition of LptDE structures shows that RBP_Oeko_ could not bind to the crystallised, “resting state” LptDE complex due to extensive overlaps that would result in several LptD and LptE (Fig. 2E). Therefore, RBP_Oeko_ binds to a SfLptDE state with a substantially different conformation compared to hitherto determined experimental structures, which are very similar to AF predictions (Fig. 1E).

Strikingly, strong density for an extra β-strand peptide is visible between LptD strands S1 and S26, *i.e*. it is bound in the lateral gate (Fig. 2B, C). Based on distinctive density indicating a Trp at the extracellular OM interface (Fig. 2B), we assigned the peptide as LptD residues Glu197-Pro209, corresponding to βN10 of the jellyroll domain (Fig. S5), and named it S0. The peptide is oriented parallel to S26 and forms a total of 14 hydrogen bonds with strands S1 and S26, with all but one of them between backbone peptide bonds. The presence of the peptide means that RBP-bound LptD has a 27-stranded barrel. The distance between the backbone atoms of strands S1 and S26 is ∼10 Å, which would likely be large enough to allow lateral passage of an LPS molecule into the OM. We therefore propose that RBP_Oeko_ has bound to an open, active state of the LptDE translocon. Molecular dynamics (MD) simulations in a model *E. coli* OM showed that all four components of the complex are stable (Fig. S6). Interestingly, removal of the bound S0 strand (while keeping RBP_Oeko_ bound) resulted in a very rapid closing of the lateral gate and the formation of several stable inter-strand hydrogen bonds (Fig. S7), suggesting that the barrel has a high plasticity.

To exclude the possibility that the formation of the LptDE-RBP_Oeko_ complex was a one-off adventitious event, we generated another cryo-EM sample using different LptDE and RBP_Oeko_ preparations. Data collection and processing (dataset 2; Table S1 and Fig. S8) resulted in an identical structure of the LptDE-RBP_Oeko_ complex (Cα r.m.s.d 0.4 Å), including the bound S0 strand. Interestingly however, when contoured at low levels, this map shows continuous density connecting the S0 lateral gate peptide to LptD strand S1 (Fig. S9 and Movie S2 and S3), confirming the assignment of strand S0 as residues 197-209 of the N-terminal LptD jellyroll domain.

According to the consensus model, LPS insertion into the OM requires not only opening of the lateral gate for diffusion of the lipid A acyl chains but also a channel to the extracellular side to allow passage of LPS polar groups such as core oligosaccharides and the O-antigen. As may be expected from the insertion of an additional β-strand, the “laterally open” LptD barrel is much wider at the front of the complex (Fig. 3A), with strands β1-β12 and EL1-EL5 displaced outwards by up to 7 Å. As a result, the pronounced dent in the closed barrel between strands 5-10 has disappeared in the open state, and the barrel is almost rectangular in shape (Fig. 3B and Movie S3). The strands and EL6-EL9 at the back of the complex are virtually identical, but, moving towards the front, the (outward) backbone shifts become again pronounced and reach 6 Å for EL11-13 and the strands connecting them. Overall, the Cα RMS values between the open and closed state are 2.7 Å for LptD and 1.2 Å for LptE. The dramatic structural changes between both states are illustrated by the backbone hydrogen bond between Thr351 in LptD EL4 and Thr95 of LptE in the closed state. In the open state, the Cα atoms of these residues are ∼18 Å apart. As another example, the hydroxyl of Ser350 in EL4 makes a hydrogen bond with the side chain of Lys735 in β25/EL13 in closed LptD but is separated by ∼19 Å in the open state, with Ser350 now hydrogen bonding to Gln299 of RBP_Oeko_. The Cα atoms of Asn345 in EL4 and Ser775 close to the LptD C-terminus move from ∼10 Å in the closed state to ∼20 Å in the open state (Fig. 3B). As a result of the conformational changes, the open state has a wide channel connecting the periplasmic space to the extracellular environment with a minimum diameter of ∼ 10 Å (Fig. 3C), wide enough to allow passage of hydrophilic LPS moieties. No such channel is present in the partially open structure of *Neisseria gonorrhoeae* LptDE and in closed LptDE complexes (Fig. 3D) (17, 19, 20, 24, 26) and the minimum pore diameters of the latter are likely too small to allow significant water passage.

**Figure 3.**
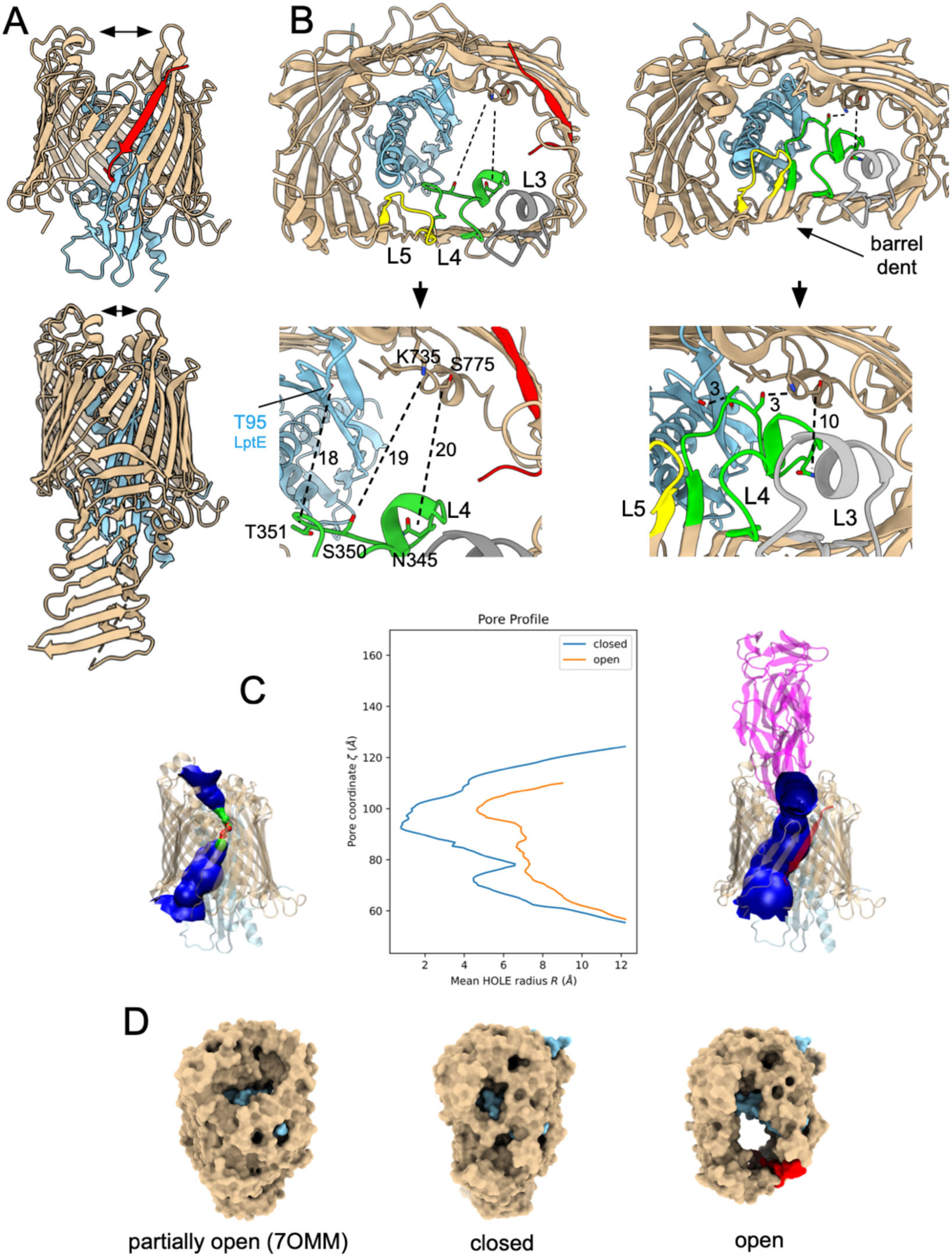
An open state of the LptDE translocon. (A) Front views of open (left panel) and closed LptDE. The lateral gate peptide is in red. RBP_Oeko_ is not shown in the left panel for clarity. The arrows indicate the distance between loops on both sides of the barrel. (B) Extracellular views of open (left panel) and closed SfLptDE. EL3 (Tyr291-Trp311), EL4 (Lys333-Thr356) and EL5 (Phe377-Ser389) are coloured grey, lime green and yellow respectively. Dashed lines indicate distances discussed in the text. The bottom panels show zoomed-in views of the same area, with distances given in Å. Views were generated from LptD superpositions. (C) HOLE (31) surface representations of resting state LptDE (PDB 4Q35; jellyroll domain not shown) on the left and SfLptDE-Oeko_RBP_ on the right. Channel surfaces connecting the periplasmic and extracellular space are shown in blue (green, diameter < 3 Å; red, diameter < 1.5 Å). The structural models are approximately aligned to the pore profiles. (D) Extracellular surfaces for (left panel) partially open NgLptDE (PDB ID 7OMM; (24)) and the closed and open states of SfLptDE. Views generated from LptD superpositions.

Besides the main particle class for the LptDE-RBP_Oeko_ complex, dataset 1 contains a smaller class of LptDE dimers for which we could generate maps to ∼3.4 Å (Fig. S4 and Table S1). Interestingly, only one of the LptDE complexes has bound RBP_Oeko_ and represents the open state, whereas the other lacks the RBP and is closed. No particle classes were observed for dimeric open or closed complexes. The LptD barrels are stacked front-to-front, with complex formation likely driven by an intermolecular disulphide bond between the Cys725 residues of both LptD protomers (Fig. 2B and Fig. S10). Given that the N-terminal jellyroll domains would clash with each other this dimeric complex is clearly non-physiological, but it allows a good appreciation of the differences between the open and closed states of the LptDE complex. Neither the open nor the closed state shows any density for LptM, presumably due to the lack of the jellyroll domain.

### Spontaneous LptD mutants abolish *E. coli* infection by Oekolampad

Maffei et al. identified two mutations in LptD that conferred resistance to a panel of LptD-targeting siphophages, including Oekolampad (Bas18). The first of these was a 4-residue deletion (Leu394-Asn397) in strand β10 that was replaced by a tyrosine residue, and the second was a 22-residue deletion (Arg657-Tyr678) in EL11 replaced by a histidine. The effect of the first deletion is hard to predict given its location within the OM-embedded part of the barrel, but it most likely affects EL5 which interacts with RBP (Fig. 4). The effect of the second mutation is easier to rationalise, given the removal of virtually the entire RBP-interacting EL11. Thus, for both Maffei mutants the generation of phage resistance is rationalised by the LptDE-RBP_Oeko_ structure. Notably, this EL11 deletion mutant was also recently obtained in a separate study of LptD-targeting phages, confirming its sufficiency to block infection (29).

**Figure 4.**
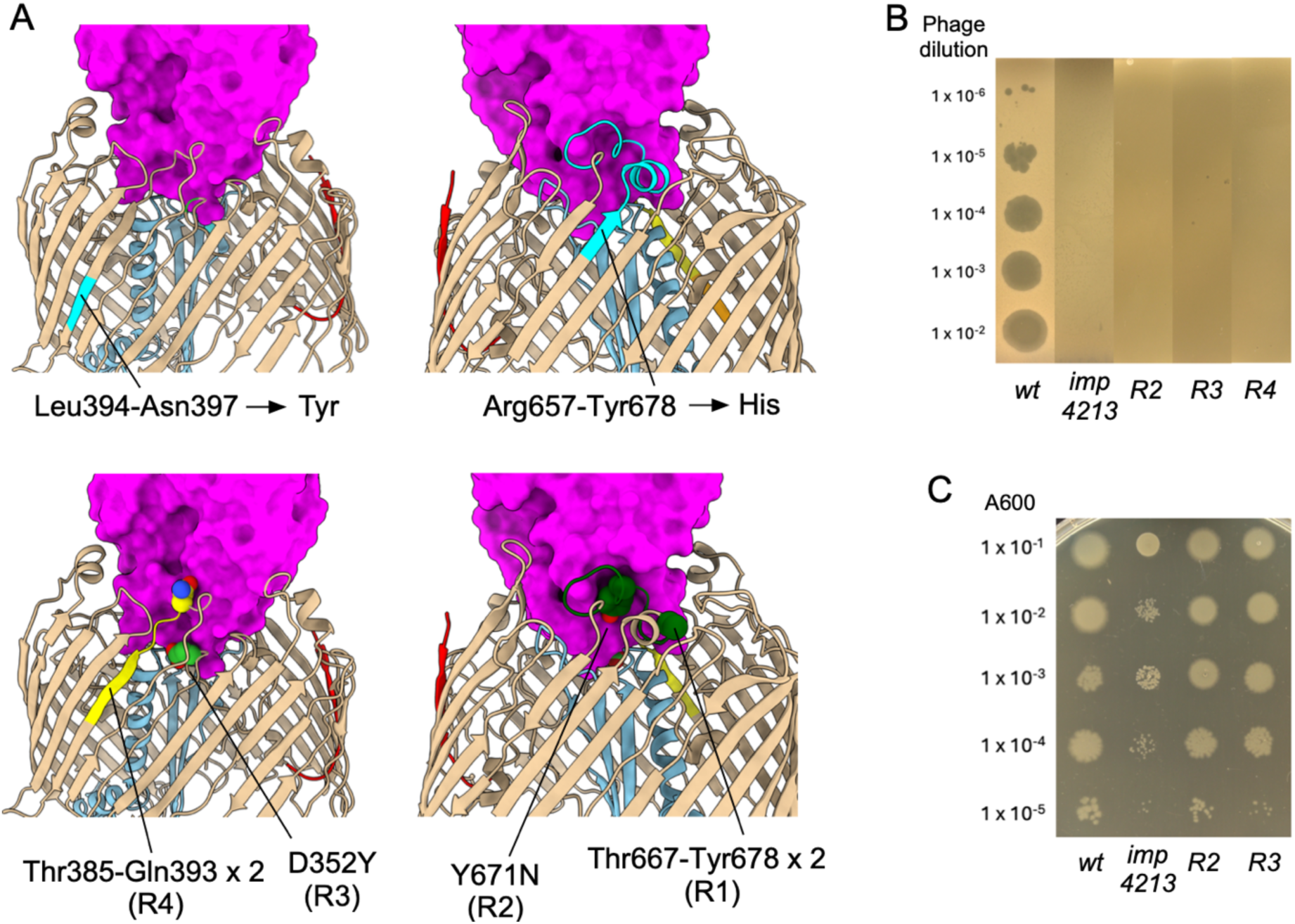
Characterisation of Oekolampad-resistant *E. coli* mutants. (A) Locations of the resistant mutants generated by Maffei et al. (top panels; mutated regions shown in cyan) and in the current study. Point mutants and duplication insertion sites are shown in space filling models, and the RBP surface is coloured magenta. Mutated EL4 residues are coloured lime green, EL5 residues yellow and EL11 residues dark green. (B) Oekolampad phage dilutions spotted on soft agar LB plates with *imp4213* and Bl21 (DE3) strains. (C) Strain dilutions spotted on LB plates with 0.5% SDS and 0.5 mM EDTA.

We were interested to further define the “resistance-space” of LptD and to this end characterised four additional resistant mutant *E. coli* Bl21(DE3) strains that spontaneously grew on Oekolampad-infected soft agar plates (Methods). Gratifyingly, all four strains had changes in their LptD as determined via whole-genome sequencing. Two mutants (R1 and R4) had insertions that both consisted of duplications. In R1, the duplicated sequence Thr667-Tyr678 in EL11 (12 residues) has inserted at Ala666, and in R4 the 9-residue sequence Thr385-Gln393 in EL5 has inserted at Asn384 (Fig. 4). Strikingly, the other two mutants (R2 and R3) are LptD point mutations: Y671N in EL11 (R2) and D352Y in EL4 (R3). All four mutants are clearly resistant to Oekolampad infection on agar plates (Fig. 4B). Intriguingly, similar mutations have recently been identified in unrelated LptD-targeting phage systems. A study of resistance evolution in *E. coli* exposed to the JNUWD phage revealed three LptD point mutations, Y671D, D226Y, and D352Y, that conferred phage resistance (32). Notably, two of these mutants (Y671D and D352Y) concern the same residue as our R2 and R3 mutants (and R3 is identical). These independent findings emphasize the key role of EL11 and EL4 in phage binding and suggest that certain sites within LptD act as evolutionary “hotspots” for resistance.

Together with the Maffei variants, these data show that resistance is readily generated via mutation of the phage receptor, even for an essential OMP complex. Given the large LptDE-RBP_Oeko_ interface and the high number of intermolecular hydrogen bonds, the generation of resistance by just a single point mutation is remarkable. To confirm this result *in vitro* and to ascertain whether the resistance is due to a defect in RBP_Oeko_ binding to LptDE, we generated the R2 and R3 point mutants in the pBAD22 expression vector and purified the LptDE complexes. Following incubation with RBP_Oeko_ as done for wild type LptDE, no co-purifying RBP_Oeko_ was observed for both mutants (Fig. 2A and Fig. S11). These results show that both point mutants abolish or weaken the high-affinity interaction of RBP_Oeko_ with LptDE and suggest this causes the observed resistance against infection. Interestingly, the *imp4213* strain (lacking the entire EL4; residues 330-352; (23)) is also resistant to infection, demonstrating that specific interactions of RBP_Oeko_ with EL4 are required for complex formation. To determine whether the resistance mutations affected LptDE function, we determined the sensitivity of the point mutant strains R2 and R3 towards SDS/EDTA in solid media. Both mutants behaved like the wild type whereas the *imp4213* control strain was, as expected, sensitive to OM stress (Fig. 4C). these findings are consistent with EL11 deletion (29), which conferred phage resistance without detectable effects on bacterial growth or recombinant protein expression. We conclude that LptDE can readily mediate resistance towards binding by small siphophages without a major loss of fitness in rich medium, indicating that phage steering via LptDE may be challenging, at least for these phages.

### Mechanism of superinfection exclusion of LptDE-targeting small siphophages

Bacteriophages produce superinfection exclusion (SE) factors that prevent non-productive phage absorption to already infected cells and cell fragments produced post-lysis. With phage adsorption to (outer) membrane receptors, this amounts to an effect by the SE factor that prevents RBP binding to the receptor. For the TonB-dependent transporter (TBDT) FhuA, the OM receptor for coliphage T5, the SE lipoprotein Llp binds to the periplasmic face of FhuA and induces “reverse” allosteric conformational changes in extracellular loops that abolish binding of the T5 RBP (3). However, since LptDE is not a TBDT it is not clear how periplasmic SE lipoproteins could abolish RBP binding to LptD. To answer this question, we initially tried to express the Oekolampad SE lipoprotein (bas18_0026; SE_Oeko_) in *E. coli* but failed. Good expression was obtained for the SE lipoprotein Rtp45 from phage Rtp (UniProt ID: Q333D9), which has 35% sequence identity to SE_Oeko_ (Fig. S3). We generated the SfLptDE-Rtp45 complex via co-overexpression in *E. coli* (Fig. 5A). The cryo-EM data contain only one major particle class, with bound Rtp45 (Table S1 and Fig. S12). The small lipoprotein (∼6 kDa) is bound to the periplasmic, left-hand side of LptDE close to β-strands 5-8 (Fig. 5B, C). Except for the N-terminal three residues including the lipid anchor, the entire protein is visible in the map. PISA analysis shows that the LptD-Rtp45 interface has an area of almost 1000 Å^2^ (CSS = 1.00). There are 13 hydrogen bonds between Rtp45 and LptD and two salt bridges. A β-sheet is present between Rtp45 residues 30-35 and LptD residues 318-323 at the base of strand β6. In addition, Rtp45 forms three hydrogen bonds with LptE on one side and two hydrogen bonds with LptM on the other, meaning it is sandwiched between both lipoproteins (Fig. 5B). Only the first ∼10 residues of LptM are visible in this map, similar to other LptDEM structures (Fig. 5B, C). The fact that LptM is bound in the presence of Rtp45 must mean that the remainder of LptM (*i.e.* the part not visible in the map) is in the periplasmic space and supports the notion that LptM might be permanently bound to the LptDE translocon via its N-terminal ∼10 residues. Besides a disulphide bond between Cys30 and Cys38, Rtp45 has a long, partially β-stranded hairpin that protrudes upwards through the LptD lumen (Fig. 5D). As is common for phage proteins, being under-represented in structural databases, the AF3 prediction of Rtp45 is very different from the experimental structure, although the Cys30-Cys38 disulphide is correctly predicted (Fig. S3). Interestingly, the model of SE_Oeko_ is more similar to the Rtp45 experimental structure, even though the predicted disulphide is incorrect as it involves the N-terminal cysteine. Despite this, it seems a reasonable assumption that the interaction of SE_Oeko_ with LptD is similar to that of Rtp45. The AF3 predictions for the LptDE-SE_Oeko_ and LptDE-Rtp45 complexes all show the SE proteins interacting with the LptD jellyroll domain (Fig. S13) and are clearly incorrect.

**Figure 5.**
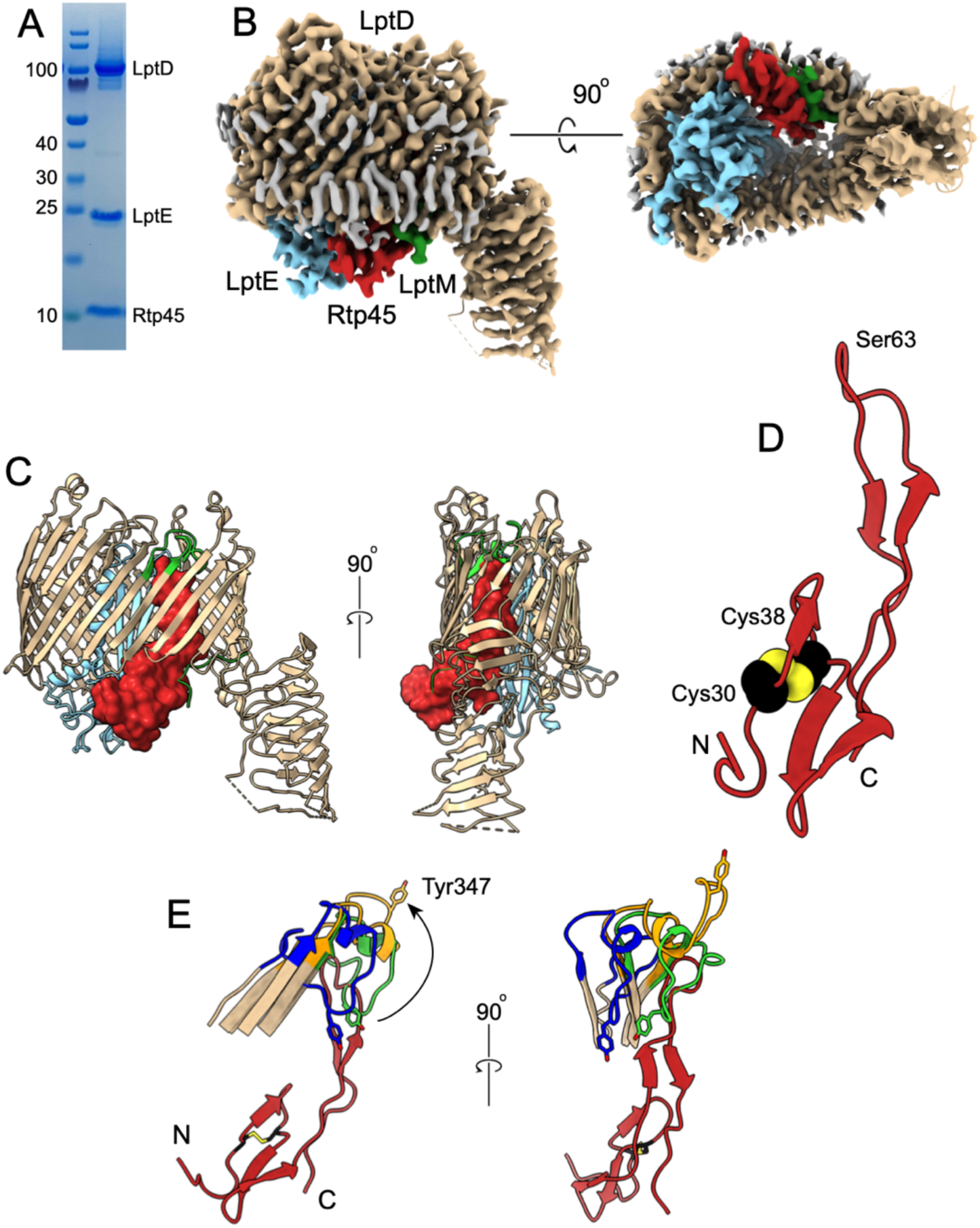
Superinfection exclusion mechanism of phages Oekolampad and Rtp. (A) SDS-PAGE gel of the SfLptDE-Rtp45 complex following SEC. (B) Cryo-EM density maps of the SfLptDE-Rtp45 complex viewed from the left-hand side and from the periplasmic space. Rtp45 is coloured maroon and is sandwiched between LptE and LptM. (C) Cartoon viewed from the left-hand side and from the front, with Rtp45 shown in surface representation. LptD EL4 is lime green. (D) Cartoon model of Rtp45 viewed in the same orientation as in C (left-hand panel), with Cys30 and Cys38 shown as space-filling models. (E) Cartoon models viewed from the left-hand side and from the front (right panel) of EL4 in LptDEM (lime green), LptDE-RBP (open state, blue) and LptDEM-Rtp45 (orange, with Rtp45 in maroon). The movement of Tyr347 in EL4 from the closed LptDEM state to the Rtp45-bound state is indicated with the arrow. The reverse movement would be blocked by Rtp45, providing a rationale for abolishing RBP binding. Views generated from LptD superpositions.

The main consequence of Rtp45 binding is that EL4 of LptD is displaced towards the extracellular side by up to 20 Å (for the Tyr347 Cα atom; Fig. 5D). This conformation of EL4, together with the presence of bound Rtp45 will cause superinfection exclusion due to (i) induction of a LptD conformation that is incompatible with RBP binding and (ii) blocking the movement of EL4 in the open state (Fig. 5E and Movie S4). Thus, the conformational changes in LptD that block binding of the phage RBP result from physical displacement of extracellular loops by the SE protein, and not from long-range allostery as in FhuA.

## Discussion

The most important and surprising finding of our work is that the RBP of the small siphophage Oekolampad, and most likely related phages, binds to an open state of the LptDE translocon, enabling experimental visualisation. The formation of the complex is extremely slow, suggesting that LptDE states enabling RBP_Oeko_ binding are very rare and/or short-lived *in vitro*, which is consistent with existing structural data for LptDE. The fact that MD simulations show that the lateral gate in LptDE-RBP_Oeko_ rapidly closes upon removal of the bound jellyroll-derived S0 segment *in silico* suggests the barrel is flexible and raises the possibility that bound RBP would not interfere with LPS translocation. Interestingly, very low levels of complex appear to be formed between LptDE lacking the jellyroll domain (LptD_Tr_E) and RBP_Oeko_ (Fig. S11), suggesting that S0 binding favours complex formation, presumably by stabilising the open state, but is not absolutely required.

The degradation of the LptD jellyroll domain required for formation of the open, RBP binding-competent LptDE state *in vitro* are likely consequences of the detergent extraction and purification procedures. However, the high specificities of receptor-RBP interactions suggest that the visualised open state is relevant *in vivo*. Indeed, translocating LPS molecules *in vivo* should induce a state of LptDE with similar properties as the one in our structures, *i.e*. with an open lateral gate and a large channel connecting the periplasmic space to the extracellular side. We therefore propose that Oekolampad and similar phages target an active state of the LptDE translocon, perhaps to ensure efficient phage replication via selecting actively growing cells. An interesting implication of the bound lateral gate peptide is that it suggests that LptDE function might be inhibited by peptidomimetics that bind stably to the open lateral gate, such as those discovered for the BamA component of the BAM complex (33, 34). To begin to test this hypothesis we synthesised the S0 peptide (LptD residues 196-207) and determined MIC values against a panel of *E. coli* strains, most of which have compromised OM barriers.

Intriguingly, while no MICs were observed for most of the strains, both the L– and D-form of the S0 peptide generated weak MICs against the *imp4213* strain (Table S2), suggesting that the absence of the LptD EL4 in this strain improves the accessibility of the lateral gate from the outside of the cell and that the S0 interaction mainly involves backbone interactions. However, it is worth noting that this mutant is generally more susceptible to non-specific, membrane-disrupting antimicrobial peptides, which may contribute to the observed activity. Whether or not effective antibiotics targeting the LptD lateral gate can be designed remains an open question; thus far, only peptidomimetics that target periplasmic parts of *Pseudomonas aeruginosa* LptDE have been obtained (35, 36). However, the availability of an open state that is likely to be populated *in vivo* may open up opportunities for *in silico* drug discovery of LptD inhibitors.

The presence of LPS-mimicking LptM within the purified LptDE complex is very intriguing. The density for the hitherto unobserved C-terminal half of LptM is weaker than that for LptDE, suggesting mobility or unresolved alternate conformations. However, the density for the first N-terminal LptM residues is comparable to that of the neighbouring LptD barrel, suggesting LptM is bound to at least the majority of LptDE complexes within the sample. Given that LptDE doesn’t copurify with LPS, it is unlikely that LptM binds to LptDE during detergent extraction. LptM must therefore be present within fully matured, functional LptDE complexes in the OM. Based on our structure, LptDE-bound LptM might block LPS transport, raising the question of how the presence of bound LptM is compatible with normal cell growth. This question is very relevant given that LptM is present in a serendipitously isolated hybrid complex between overexpressed, His-tagged SfLptE and endogenous EcLptD (see Methods). Thus, the observed LptD-LptM interaction is not an artefact of LptD overexpression. Some insights into this conundrum may be gained from a quantitation of the *E. coli* proteome (37), which showed that LptD and LptE are present at ∼400-600 copies per cell depending on the strain and growth media. By contrast, the ABC transporter components LptC, LptF and LptG are much less abundant with ∼60-120 copies per cell, whereas LptM is very abundant (∼2000-2500 copies/cell). Thus, based on these numbers, only a minor fraction of LptDE molecules can be active at any time, *i.e*. physically engaged within a periplasmic protein bridge. If, following biogenesis, all LptDE complexes contain LptM and the intermembrane Lpt protein bridges are very stable, it is possible that LptM is cleared from its binding site by translocating LPS molecules in only a small fraction of the LptDE pool, which might be hard to detect within a cryo-EM dataset. An interesting implication of this hypothesis, supported by a recent study (26), is that LptDE may mediate surface exposure not only of LPS but also of LptM and perhaps other small lipoproteins like its homologue in *Borrelia burgdorferi* (38). Whether this has functional significance is not clear, but it would explain, for example, why part of the Braun’s lipoprotein (Lpp) pool is surface exposed (39). Alternatively, our data might mean that LptM is an integral component of the LptDE translocon, perhaps to assist LPS translocation, a notion supported by recent data that suggest LptM binding weakens interactions between S1 and S26 of the lateral gate (25). Although our LptDEM structure suggests that the part of LptM that is present in the LptDE lumen would block the LPS translocation pathway, this blockage could be relieved by movement of this part of LptM into the periplasmic space while remaining bound to the translocon via its N-terminus, as observed in the LptDE-Rtp45 complex (Fig. 5) and other recent structures (25, 26).

## Materials and Methods

### Cloning and Mutagenesis

The pBAD22 plasmid containing *Shigella flexneri* LptD and His6-tagged LptE (17) was generously provided by the Huang lab (Institute of Biophysics, Chinese Academy of Sciences). Truncation of LptD (Δ26-201) and the insertion of a TEV protease recognition site immediately upstream of the C-terminal His-tag on LptE were performed using site-directed mutagenesis with the Q5 High-Fidelity DNA polymerase kit (New England Biolabs), following the manufacturer’s protocol. Mutagenic primers were designed using the NEBaseChanger tool. The full-length genes encoding for Rtp44, RBP_Oeko_, RTP45 and SE_Oeko_ were synthesised by Eurofins Genomics, and cloned into the pET28b expression vector (Novagen) via *Nco*I and *Xho*I sites, appending the sequence LEHHHHHH to the proteins. Genomic DNA was isolated from spontaneous phage resistant *E. coli lptD* mutants R1-R4 via the Sigma-Aldrich GenElute bacterial gDNA kit and sent for whole genome sequencing (Eurofins genomics). The *lptD* genes from the resistant mutants were amplified via PCR using primers that incorporated *Nco*I and *Hind*III restriction sites at the 5’ and 3’ ends, respectively. Both amplified insert and SfLptDE-pBAD22 vector were digested with *Nco*I and *Hind*III (New England Biolabs), PCR purified and ligated using T4 DNA ligase (New England Biolabs). The mutants are the following: R1, Thr667-Tyr678 duplication at Ala666; R2, Y671N; R3, D352Y; R4, Thr385-Gln393 duplication at Asn384.

### Protein Expression and Purification

#### LptDE(M) and mutants

Full-length (LptDE), truncated (LptD_Tr_E), and mutant variants (*imp4213*, *R2* and *R3*) of LptDE were moved into electrocompetent *E.coli* BL21(DE3)Δcyo cells (ΔcyoB, truncated *cyoA* and *cyoC*) via electroporation. Overnight starter cultures were grown in LB medium supplemented with ampicillin (100 μg/mL) and used to inoculate 1 L flasks of LB (100 μg/mL ampicillin) to a starting OD_600_ of ∼0.05. Cultures were grown at 37 °C, 180 rpm until OD_600_ reached 0.8–1.0, at which point expression was induced with 0.1% (w/v) arabinose. Full-length LptDE and its mutants were expressed at 37 °C, 180 rpm for 2.5 hours post-induction, whereas LptD_Tr_E was expressed at 18 °C and 150 rpm for 20 hours post-induction. For co-expression of LptDE and Rtp45, cells were sequentially transformed with SfLptDE-pBAD22 and Rtp45-pET28. Co-expression was performed under the same conditions as full-length LptDE; no IPTG was added to express Rtp45, as leaky expression from the T7 promoter was sufficient to produce excess Rtp45 relative to LptDE.

Cells were harvested by centrifugation at 4 °C for 20 minutes at 4,200 rpm (Beckman J6-HC, JS-4.2 rotor), and resuspended in TBS (20 mM Tris-HCl, pH 8.0, 300 mM NaCl) supplemented with DNase I. Cells were resuspended with a Dounce homogenizer, followed by two passes through a Constant Systems cell disruptor (0.75 kW model) at 20,000 psi. Lysates were clarified by ultracentrifugation (42,000 rpm, 50 minutes, 4 °C; Beckman Optima XE-90, 45Ti rotor), and the membrane pellet was resuspended in TBS containing 2% (w/v) LDAO. After homogenization and stirring for 1 hour at 4°C, insoluble material was removed by ultracentrifugation (30,000 rpm, 30 minutes, 4°C; Beckman Optima XE-90, 45Ti rotor). The supernatant containing solubilized LptDE was loaded onto a gravity-flow IMAC column packed with Ni²⁺-charged Chelating Sepharose (Cytiva), pre-equilibrated in wash buffer (20 mM Tris-HCl, pH 8.0, 300 mM NaCl, 30 mM imidazole) + 0.15% (w/v) DDM. The column was washed with 30 column volumes of wash buffer. Bound protein was eluted with 3 column volumes of elution buffer (20mM Tris-HCl, pH 8.0, 300mM NaCl, 200mM imidazole) + 0.15% (w/v) DDM. The eluate was concentrated via a 100 kDa MWCO Amicon Ultra centrifugal filter (Millipore) and loaded on a Superdex-200 16/600 SEC column (Cytiva), equilibrated in 10mM Hepes, 100mM NaCl and 0.03% DDM. When required (e.g. for removal of excess LptE), an analytical SEC column (Superdex 200 Increase 10/300 GL column; Cytiva) was run next, with the same buffer as above. Complexes for cryoEM were concentrated to 5-8 mg/ml. Yields of wild type and R2/R3 mutants of SfLptDE were 0.5-1 mg/l cell culture.

When we carried out the above expression and purification procedures for a pBAD22-encoded *imp4213* variant of SfLptDE, we obtained low amounts of LptD and, as usual, an excess of LptE after IMAC. Following two rounds of SEC we obtained a reasonably well-defined peak that was concentrated, and data were collected by cryoEM. Upon inspection of the maps, it became evident that, instead of the *imp4213* variant lacking EL4, we had obtained a complex between SfLptE and EcLptD (Fig. S1). Presumably, expression levels of *imp4213* were very low and/or the protein was unstable during purification. Given that EcLptD and SfLptD are virtually identical (differences are L_Ec_453F_Sf_, R_Ec_657H_Sf_, W_Ec_691R_Sf_ in LptD and T_Ec_190M_Sf_ in LptE) we did not pursue structure determination of a wild type SfLptDE complex.

TEV cleavage of SfLptDE was carried out only when necessary (*i.e*. for isolation of the LptDE-RBP_Oeko_ complex) since it consistently reduced overall protein yield and the required reducing conditions could disrupt LptD disulphide bonds. Following initial IMAC purification, samples were concentrated and diluted at least 20-fold into cleavage buffer (50mM Tris-HCl, pH 8.0, 0.5mM EDTA, 0.2mM TCEP, 0.1% (w/v) DDM). Protein concentration was determined by absorbance at 280 nm, and His-tagged TEV protease was added at a ratio of 1 mg per 5 mg of protein. The digestion was carried out overnight at 4°C. If present, precipitate containing excess LptE was removed by centrifugation, and the supernatant was adjusted to 250mM NaCl and 10mM imidazole before being applied to a second IMAC column to remove TEV protease, uncleaved LptDE, and His-tagged fragments. Flowthrough and wash fractions were concentrated and applied to a Superdex 200 Increase 10/300 GL column (Cytiva) equilibrated in 10 mM HEPES, pH 7.5, 100 mM NaCl, 0.05% (w/v) DDM. Peak fractions were pooled, concentrated, and flash-frozen in liquid nitrogen for storage at –80 °C.

#### RBP_Oeko_

RBP_Oeko_-pET28 plasmid was transformed into electrocompetent *E. coli* BL21(DE3) cells. Overnight cultures grown in LB with kanamycin (35 μg/mL) were used to inoculate 1 L flasks (35 μg/mL kanamycin, starting OD_600_ ∼0.05). Cultures were grown at 37 °C, 180 rpm, until OD_600_ reached 0.4–0.6, then chilled at 4 °C for 1 hour prior to induction with 400 μM IPTG. Expression proceeded for 20 hours at 18 °C, 150 rpm. Cells were harvested and lysed as above, and clarified by centrifugation (19,000 rpm, 30 minutes, 4 °C; Beckman JA-25.50 rotor). The supernatant was applied to a gravity-flow IMAC column packed with Ni²⁺-charged Chelating Sepharose (Cytiva) pre-equilibrated in wash buffer. After washing with 30 column volumes of wash buffer, protein was eluted with 3 column volumes of elution buffer and concentrated using a 30 kDa MWCO Amicon Ultra centrifugal filter (Millipore). The protein was further purified by SEC using a Superdex 200 Increase 10/300 GL column equilibrated in 10 mM HEPES, pH 7.5, 100 mM NaCl, 10% glycerol. Peak fractions were pooled, concentrated, and flash-frozen in liquid nitrogen for storage at –80 °C. Yields were ∼10-15 mg/l cell culture.

### DE-RBP complex formation, purification and time course analysis

To assess the formation kinetics of the DE–RBP complex, purified SfLptDE and RBP_Oeko_ were mixed at a 1:3 molar ratio (DE:RBP) in buffer containing 10 mM HEPES, pH 7.5, 100 mM NaCl, and 0.05% (w/v) DDM. The sample was incubated at 4°C, and aliquots were withdrawn at 4.5, 24, 48, 96, and 168 hours. Each timepoint sample was subjected to size-exclusion chromatography using a Superdex 200 Increase 10/300 GL column (Cytiva) equilibrated in 10 mM HEPES, pH 7.5, 100 mM NaCl, 0.05% (w/v) DDM. The leading half of the major elution peak was collected, concentrated, and analysed by SDS-PAGE. Band intensities were quantified using ImageJ software. The ratio of RBP to LptE band intensity was plotted over time to assess complex formation. Based on the time course analyses, large-scale tag-less LptDE and tagged RBP_Oeko_ incubations for cryoEM were typically purified after 48 hours of incubation. Following incubation with RBP_Oeko_ at a 3:1 molar ratio (RBP:LptDE), the sample was applied to a gravity-flow IMAC column equilibrated in wash buffer + 0.15% (w/v) DDM to remove free LptDE. The column was washed with 20 column volumes of wash buffer and bound LptDE-RBP complex and RBP were eluted with 3 column volumes of elution buffer + 0.15% (w/v) DDM. The eluate was concentrated and further purified by SEC using a Superdex 200 Increase 10/300 GL column equilibrated in 10mM HEPES, pH7.5, 100mM NaCl, 0.04% (w/v) DDM.

### CryoEM sample preparation, data collection and data processing

#### Data collection

Quantifoil R1.2/1.3 Cu 300 mesh holey carbon grids were glow-discharged in air using a PELCO easyGlow system (12mA, 30 seconds). Samples were applied to the grids (3μl), and blotting followed by plunge freezing into liquid ethane was performed using a Vitrobot Mark IV (Thermo Fisher Scientific) operated at 4°C and 100% humidity. Grids were blotted for 6, 8 and 10 seconds with a blot force of 6. Cryo-EM movies were collected on a Titan Krios microscope operating at 300 kV. Detailed data acquisition parameters are provided in Table S1.

#### Data processing and structure refinement

Movies were imported into CryoSPARC (40) for processing. Motion correction and CTF estimation were performed with micrographs with poor CTF fits or low image quality excluded. Particles were extracted with Fourier cropping to reduce data size, followed by one or two rounds of 2D classification. Ab-initio reconstruction was conducted using both selected “good” 2D classes and excluded particles (junk). One to three rounds of heterogeneous refinement were performed using both the protein and junk classes. Particles from the final protein class/es were re-extracted at full resolution and final reconstructions were obtained using non-uniform (41) and local refinement. Detailed processing workflows for each structure are provided in SI Figs. 1, 4, 8 and 12. Structural models were generated using AlphaFold (42) and fit into cryo-EM density maps using UCSF ChimeraX (43). Rigid-body refinement was then performed in PHENIX (44). Models were then manually rebuilt and corrected in Coot (45). Final real-space refinement was carried out in PHENIX.

### Phage spot assays

*E. coli* strains (BL21*Δcyo*, *imp4213*, *R2*, *R3* and *R4*) were grown in 5mL LB at 37°C with shaking. LB soft agar (0.5% w/v) was prepared and melted prior to use. The soft agar was supplemented with 20mM MgSO_4_ and 5mM CaCl_2_ and inoculated with the appropriate bacterial strain to an OD_600_ of ∼0.1, poured onto LB agar plates, and allowed to solidify at room temperature. The Oekolampad phage (bas18) was freshly diluted to 10^-2^ to 10^-6^ in 20 mM HEPES (pH 7.5), 10 mM MgCl_2_, and 10% glycerol. Once the top agar had set, 5μl of each phage dilution was spotted on the surface. Plates were allowed to absorb the spots and were then incubated at 37°C for 24 hours.

### SDS/EDTA sensitivity assays

*E. coli* strains *R2*, *R3*, *R4*, BL21*Δcyo*, and *imp4213* were compared for sensitivity to detergent and chelator stress. LB agar plates were prepared with 0.5% SDS and 0.5 mM EDTA. Overnight cultures of each strain were used to inoculate 5 mL of fresh LB and grown at 37 °C with shaking until reaching an OD_600_ of 0.4–0.6. Cultures were then diluted to an OD_600_ of 0.1 in LB. Ten-fold serial dilutions were prepared in LB, and 1 μL of each dilution was spotted onto the SDS-EDTA plates. Plates were incubated at 37 °C, and growth was assessed after 16–24 hours to evaluate detergent and chelator sensitivity across strains.

### Molecular Dynamics Simulations

The RBP structure predicted by Alphafold2(42) was taken to model the missing region from the cryo-EM density. The AF2 RBP and the cryo-EM structure of the RBP were aligned using PyMOL(46), which gave an RMSD of 0.581 Å for 223 Cα pairs out of 257. The structure was then further aligned on 5 residues either preceding or succeeding the missing region (residues 26, 112, 143, 205 and 235), giving an RMSD of 0.262 Å. Residues 1-25, 113-142, and 206-234 of the RBP were modelled using the aligned Alphafold2 structure. This region was energy minimized while the rest of the protein complex’s atoms were restrained using position restraints of 10,000 kJ/mol.

Each simulation system had three replicates, and each replicate was individually prepared using the CHARMM-GUI membrane builder (47–49). The protein complex was embedded within a model *E. coli* outer membrane, containing *E. coli* R1-LPS without O-antigen in the outer leaflet, and 1-palmitoyl 2-cis-vaccenic phosphatidylethanolamine, 1-palmitoyl 2-cis-vaccenic phosphatidylglycerol, and cardiolipin (1-palmitoyl 2-cis-vaccenic 3-palmitoyl 4-cis-vaccenic diphosphatidylglycerol) in the inner leaflet in a 90:5:5 ratio. The system was solvated with water and K^+^ Cl^-^ ions at 0.2 M. Calcium ions were used to neutralize negatively charged phosphate groups of LPS. Each simulation box was 14 nm x 14 nm x 16 nm.

Energy minimization of the systems was conducted using the steepest descent algorithm until the maximum force was below 500 kJ/mol. Systems were first equilibrated within the NVT ensemble for 200 ps, with a time step of 1 fs, using the v-rescale thermostat (50), with a time constant of 1 ps, at a temperature of 313 K. Initial velocities were assigned randomly for each replicate. Subsequent further equilibration and production runs were conducted within the NPT ensemble using a time step of 2 fs. Pressure was maintained at 1 bar using the c-rescale barostat (51) with a time constant of 5 ps. Covalent bonds involving hydrogen were constrained using the P-LINCS algorithm (52). Electrostatic interactions were described using the Particle Mesh Ewald method (53) Equilibration simulations of duration 31 ns of were performed while gradually releasing position restraints on amino acids and lipids. A further 30 ns of equilibration was performed without any position restraints. It was within this state that the removal of the bound S0 strand resulted in a very rapid closing of the lateral gate. Production runs were 2 µs. All simulations used the CHARMM36m (54) forcefield and MD simulations were performed using the GROMACS (55) software package (version 2022.4).

HOLE (31, 56) was run using the HOLE2 implementation within the MDAnalysis (57, 58). The centre of mass between Asn345 and Ile777 served as the starting point for HOLE analysis in both the closed and open system. The DSSP algorithm (59) was used to analyse secondary structure, implemented through MDTraj (60). Hydrogen bond numbers were calculated using MDAnalysis, using the backbone atoms of the S0 peptide and the surrounding beta barrel. A quite permissive Donor-H-Acceptor angle cutoff for hydrogen bonds of 135.0°, distance cutoff between donor-hydrogen pairs of 2.5 Å, and distance cutoff between donor and acceptor of 3.5 Å was used, although this falls well within the Baker-Hubbard definition (61).

### Minimum Inhibitory Concentration Assays (MICs)

MICs were carried out according to the Clinical Laboratory Standards Institute (CLSI) M100 guidelines. Bacterial cell strains were spread onto LB-agar plates and grown overnight. For *E. coli* strains: ATCC25922, *imp4213*, *ΔsurA*, *ΔwaaD*, *ΔdsbA*, *ΔdsbC*, *ΔlptM* and DC2, inoculate was prepared by suspending colonies from the plate into 0.9% saline to a cellular density corresponding to a 0.5% McFarland standard, then diluted 1:400 into either Mueller-Hinton broth (MHB, 70192) (Millipore). For hyperporinated *E. coli* strain GKCW102 and its control parent strain GKCW101, an overnight starter culture was grown in MHB then used to inoculate fresh media and grown to an OD_600_ of 0.3. FhuA pore expression was induced by the addition of arabinose to a final concentration of 0.1%, then cells were grown to an OD of 1. This culture was diluted 1:2000 and used as the MIC inoculate. 200 µl of inoculum from each strain was added to a 96-well plate containing a dilution gradient (with a top concentration of 64 µg/mL) of S0 peptide solubilised in DMSO. Both the L– and the D-form of the peptide were assayed in the same way. All cell lines were separately treated with antibiotic controls. Plates were incubated overnight at 37 °C, then checked for growth following CLSI guidelines.

## Acknowledgements

We thank Yihua Huang ((Institute of Biophysics, Chinese Academy of Sciences) for the gift of the pBAD22-*SflptDE* expression plasmid. ED acknowledges the support of the Barbour Foundation (https://www.barbour.com/uk/the-barbour-foundation) for a PhD fellowship. SK is funded by the Engineering and Physical Sciences Research Council via an Established Career Fellowship (EP/V030779) and HECBioSim (EP/R029407/2). SK and RC acknowledge allocation on ARCHER2 provided by EPSRC via HECBioSim. We thank Augustinas Silale for assistance in CryoEM data collections. SA was funded by a KTP award (10086683) from Innovate UK. Bacterial cell knockouts E. *coli* ΔSurA, ΔWaaD, ΔLptM, ΔDsbA, ΔDsbC were derived from the Horizon Discovery KEIO collection. We thank Helen Zgurskaya (The University of Oklahoma) for the kind gift of the hyperporinated *E.coli* strains GKCW101 and GKCW102, Thomas Silhavy (Princeton University) for the gift of the *imp4213* strain, and we thank the Bicycle Therapeutics chemistry team for the production of the S0 peptides.

## Author contributions

BvdB conceived and led the project, managed cryo-EM data collections, and wrote the paper. BvdB and ED generated samples, determined cryo-EM structures and performed experiments. ED assisted in writing the paper. AB maintained cryoEM computing resources and assisted in cryoEM data collection. RC performed MD simulations, supervised by SK. SA and HN performed MIC analysis.

## Data availability

Original data created for the study are or will be available in a persistent repository upon publication. CryoEM maps have been deposited in the Electron Microscopy Data Bank and atomic coordinates have been deposited in the Protein Data Bank with accession codes EMD-54169/9RPR (LptDEM), EMD-54171/9RPT (SfLptDE-RBP monomer dataset 1), EMD-54173/9RPW (SfLptDE-RBP dimer), EMD-54175/9RQI (SfLptDE-RBP monomer dataset 2), and EMD-54170/9RPS (SfLptDEM-RTP45).

## Supplementary Information

**Fig. S1.**
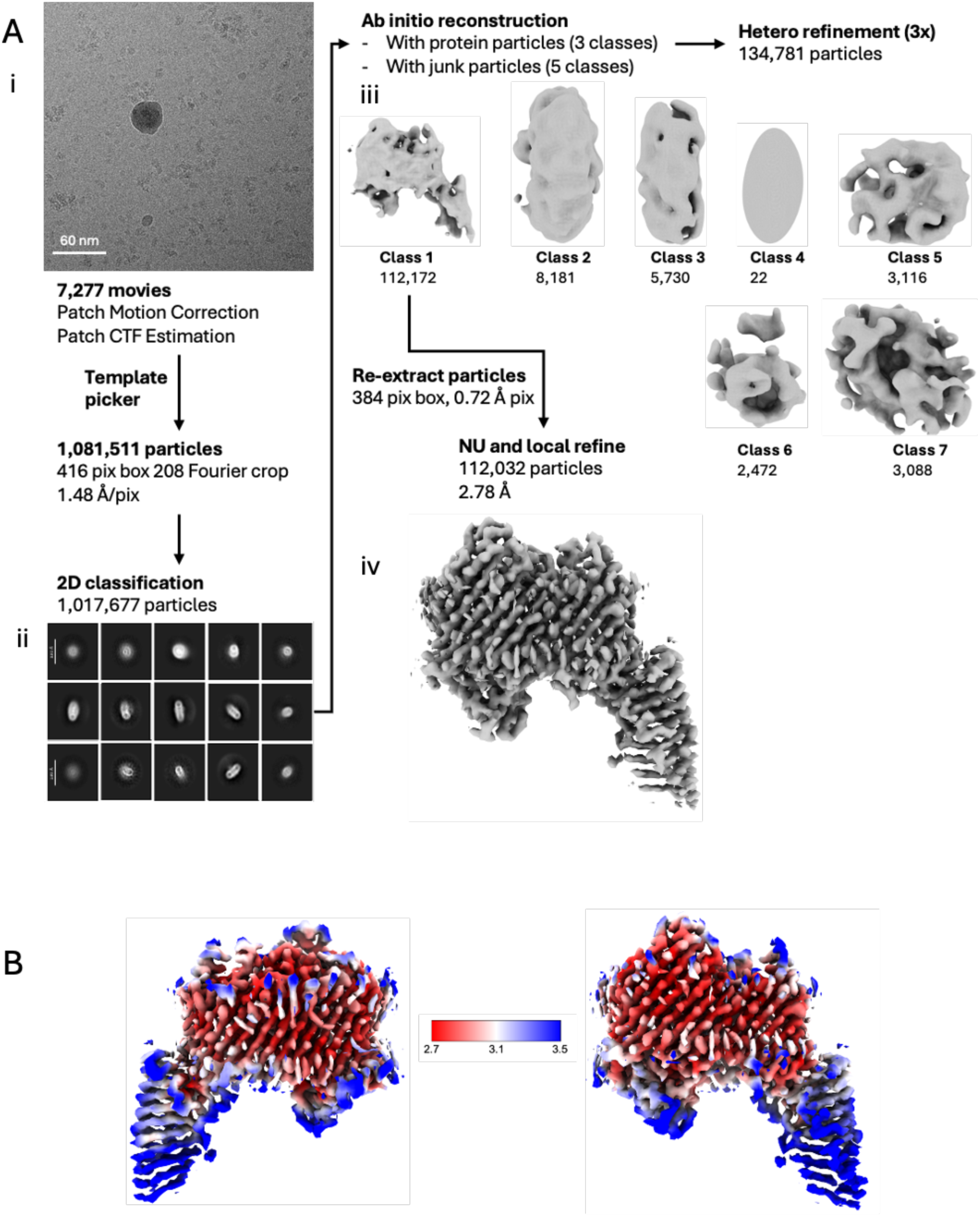
Cryo-EM data processing for *Sf*LptE-*Ec*LptD-*Ec*LptM complex. (A) Workflow summary. 7,277 movies were collected and imported into CryoSPARC (1). After motion correction and CTF estimation, low-quality micrographs were excluded. Particles were extracted with Fourier cropping and subjected to 2D classification to remove obvious junk (i, example micrograph; ii, representative 2D classes). Ab-initio reconstruction produced volumes used in three rounds of heterogenous refinement with one “good” and six “junk” classes (iii, volumes from final round). particles were re-extracted at full resolution for non-uniform (2) and local refinement, yielding a final 2.78 Å map (iv, final volume). (B) Local resolution estimation of the final map.

**Fig. S2.**
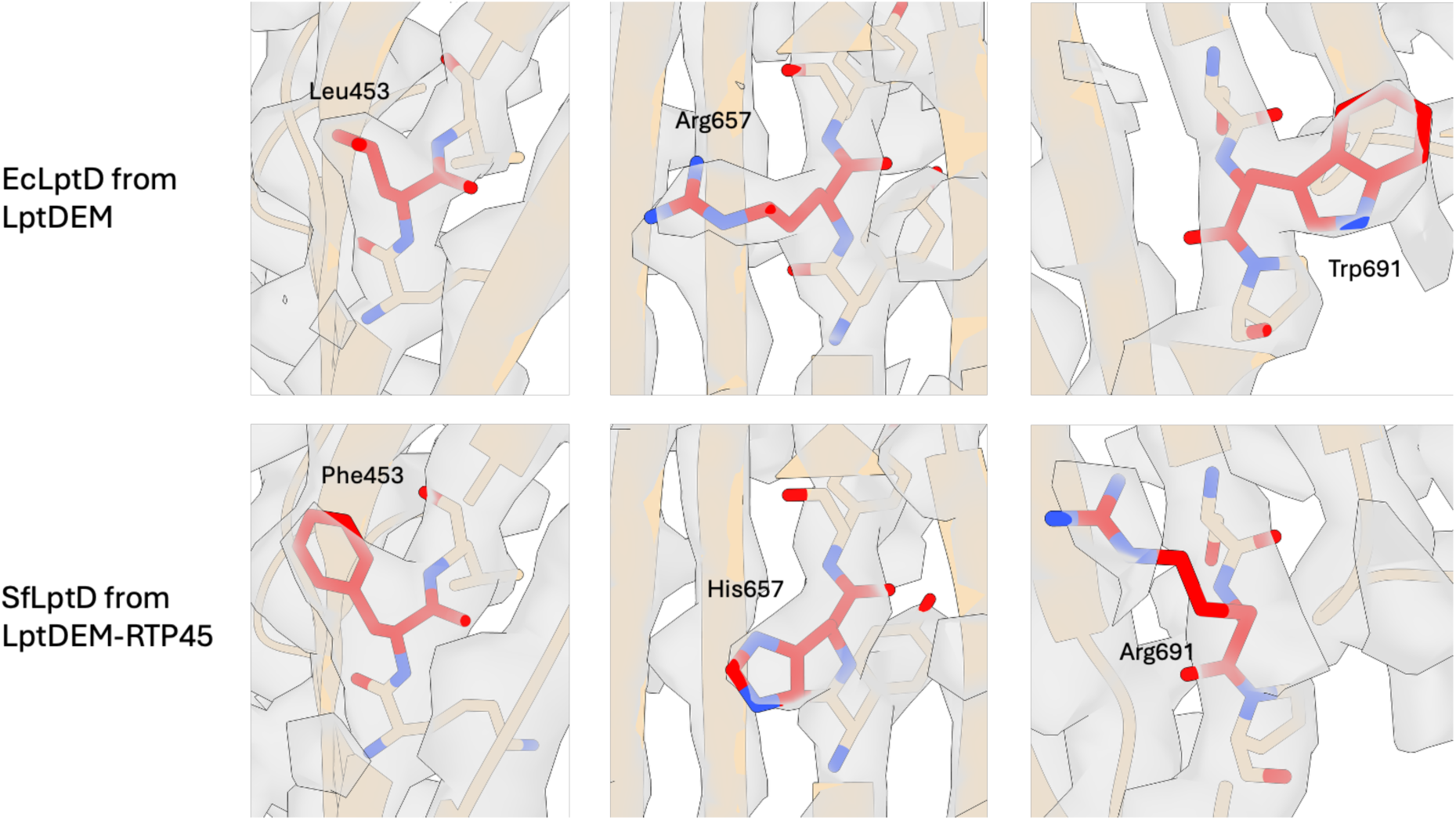
Amino acid differences between *E. coli* and *S. flexneri* LptD. The three amino acid differences at positions 453, 657 and 691 are highlighted in red. These are shown in the *Ec*LptD-*Sf*LptE-*Ec*LptM and the *Sf*LptDE-*Ec*LptM-RTP45 maps and models.

**Fig. S3.**
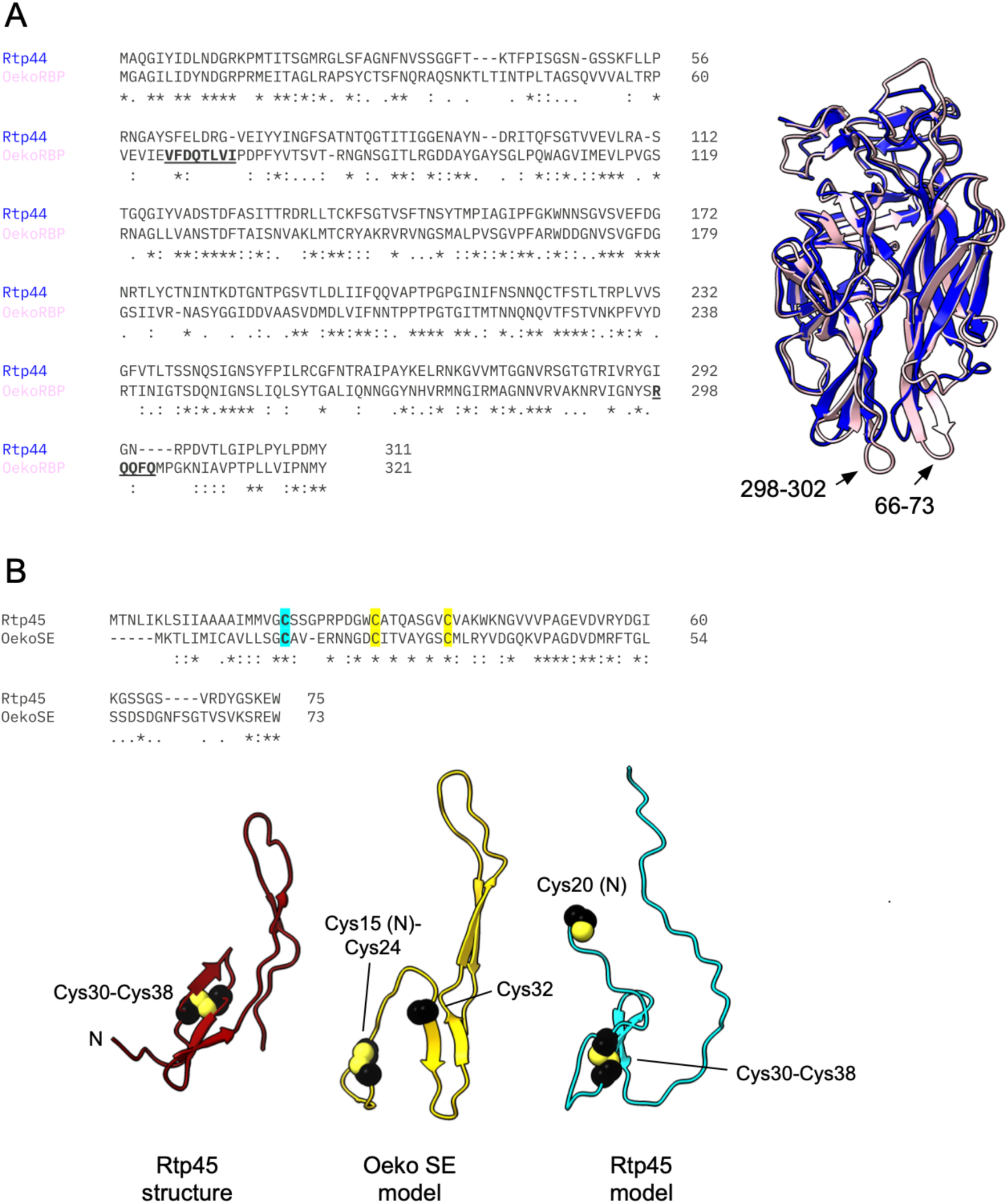
Structural comparisons of LptD-targeting RBPs and SE proteins. (A) Sequence alignment of Rtp44 and RBP_Oeko_, with a superposition of AF3-predicted models on the right. The divergent tips of two loops that likely mediate slightly different interactions with LptD are underlined and indicated. (B) Sequence alignment of Rtp45 and SE_Oeko_. The lipid anchor cysteine is indicated in cyan, and those forming the intramolecular disulphide in yellow. The bottom panel shows (from left) the experimental Rtp45 structure and representative AF3 models of SE_Oeko_ and Rtp45.

**Fig. S4.**
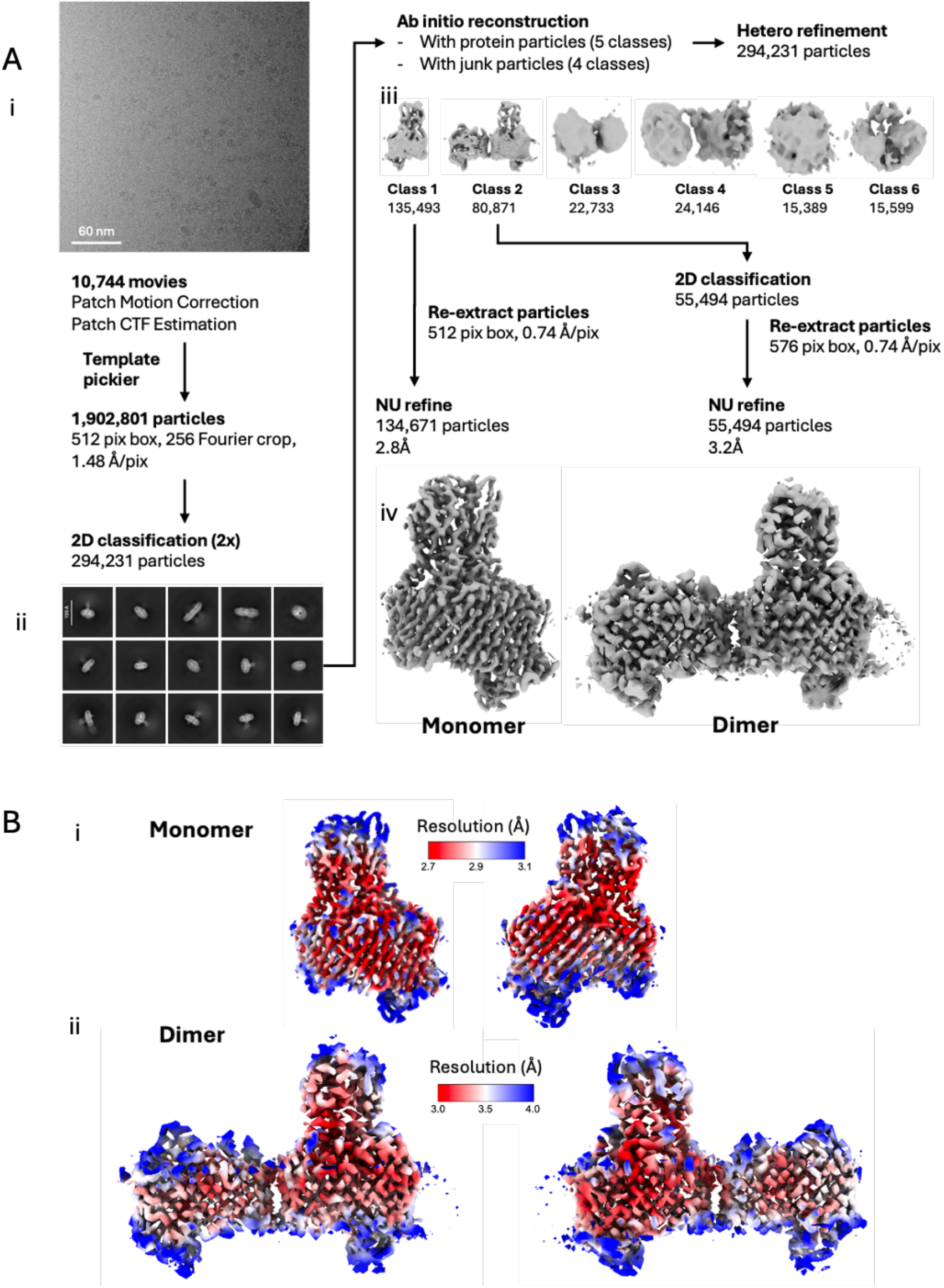
Cryo-EM data processing for the *Sf*LptD-RBP dataset 1 (monomer and dimer). (A) Workflow summary. 10,744 movies were collected and process in CryoSPARC. After motion correction and CTF estimation, low-quality micrographs were removed. Particles were extracted with Fourier cropping and subjected to two rounds of 2D classification (i, example micrograph; ii, representative 2D classes). Ab-initio reconstruction followed by heterogenous refinement using one monomer, one dimer and four junk classes (iii, volumes). Monomer and dimer particles were re-extracted at full resolution for non-uniform refinement, producing final maps of 2.8Å (monomer) and 3.1Å (dimer) (iv, final volumes). (B) Local resolution estimation of final maps: (i) monomer, (ii) dimer.

**Fig. S5.**
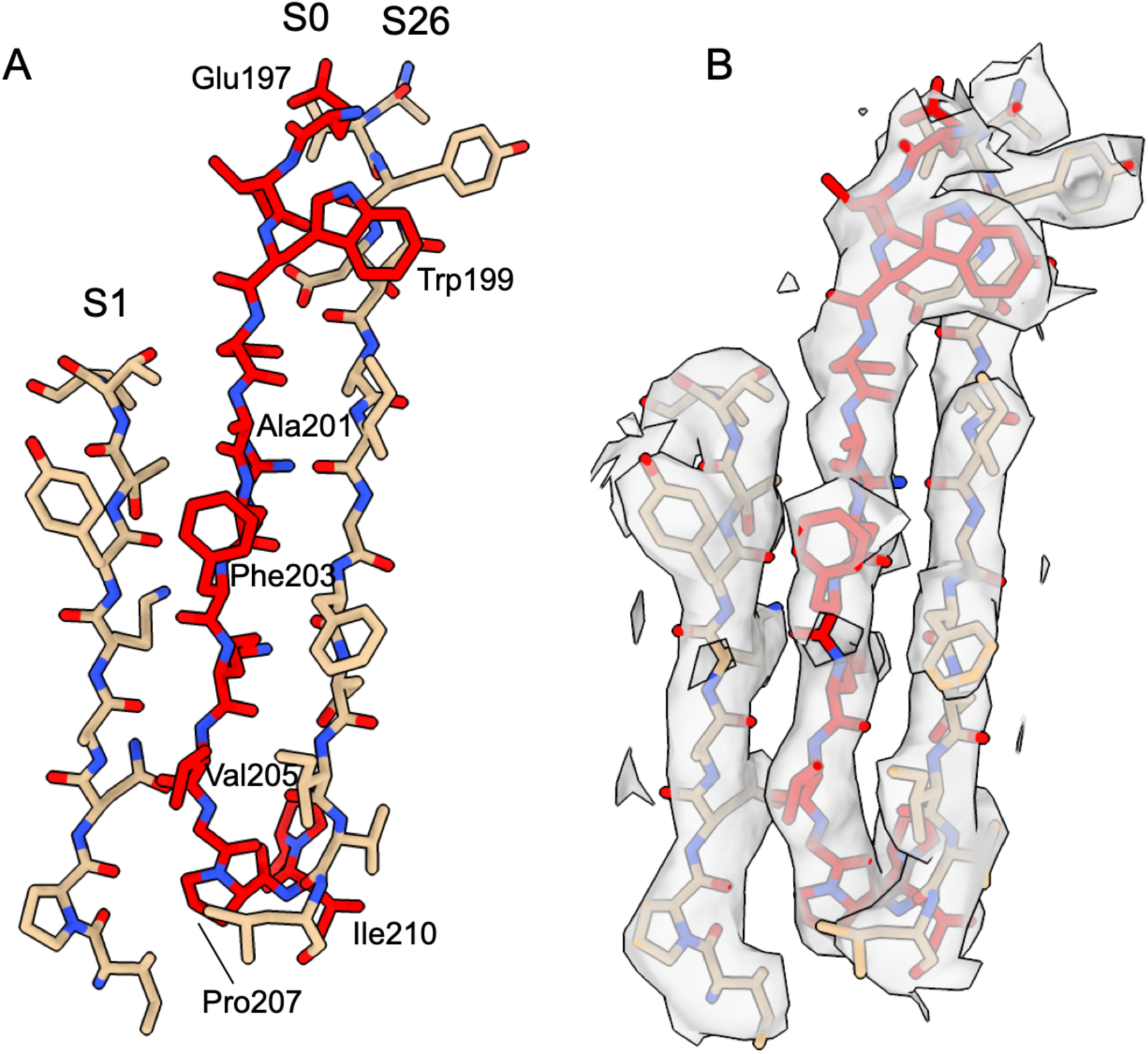
The lateral gate strand S0 from LptDE-RBP dataset 1. (A) Stick model of LptD strands S0 (red), S1 and S26. Outward facing residues in S0 have been labelled. Corresponding map for dataset 1 of the region in (A).

**Fig. S6.**
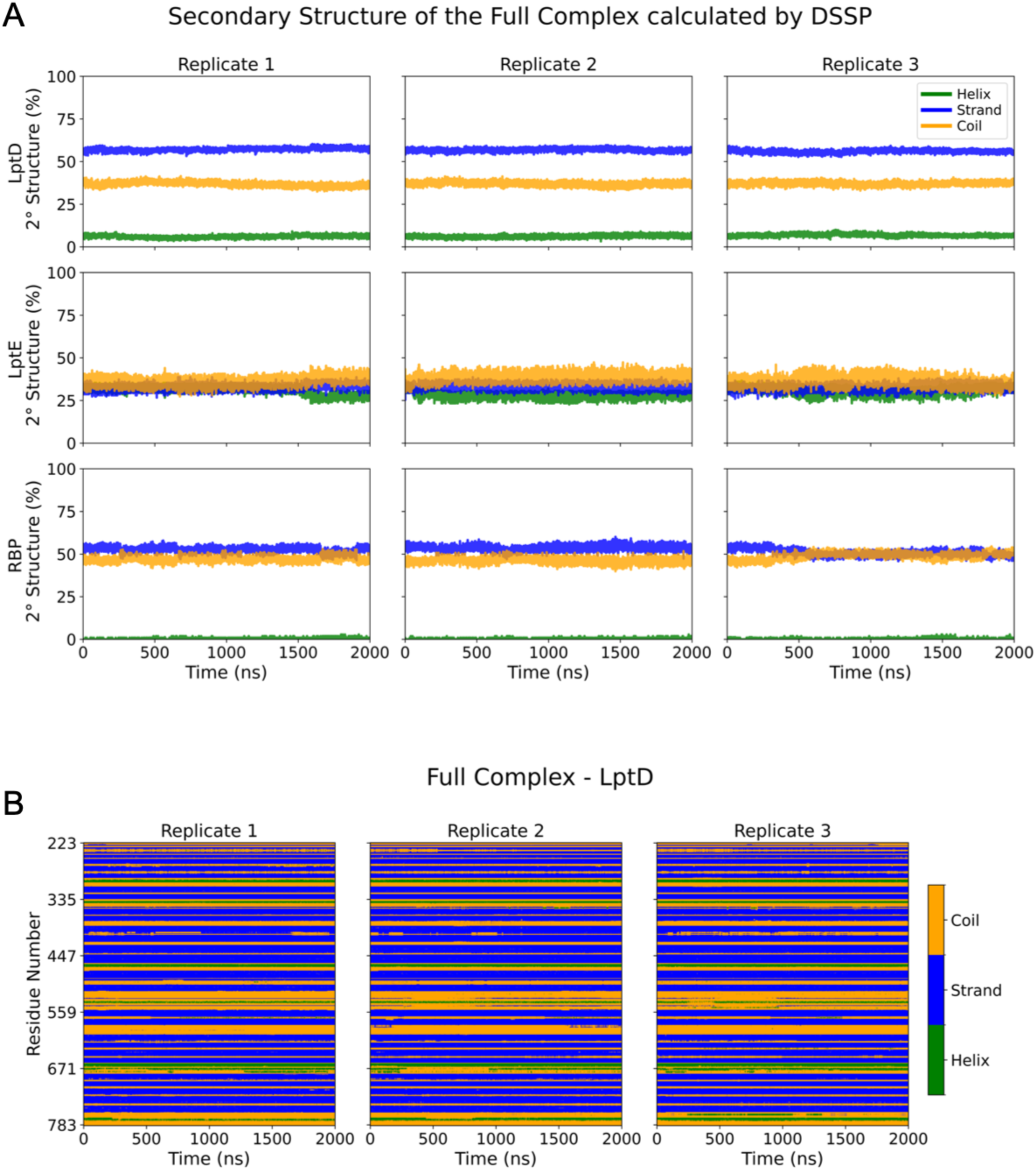
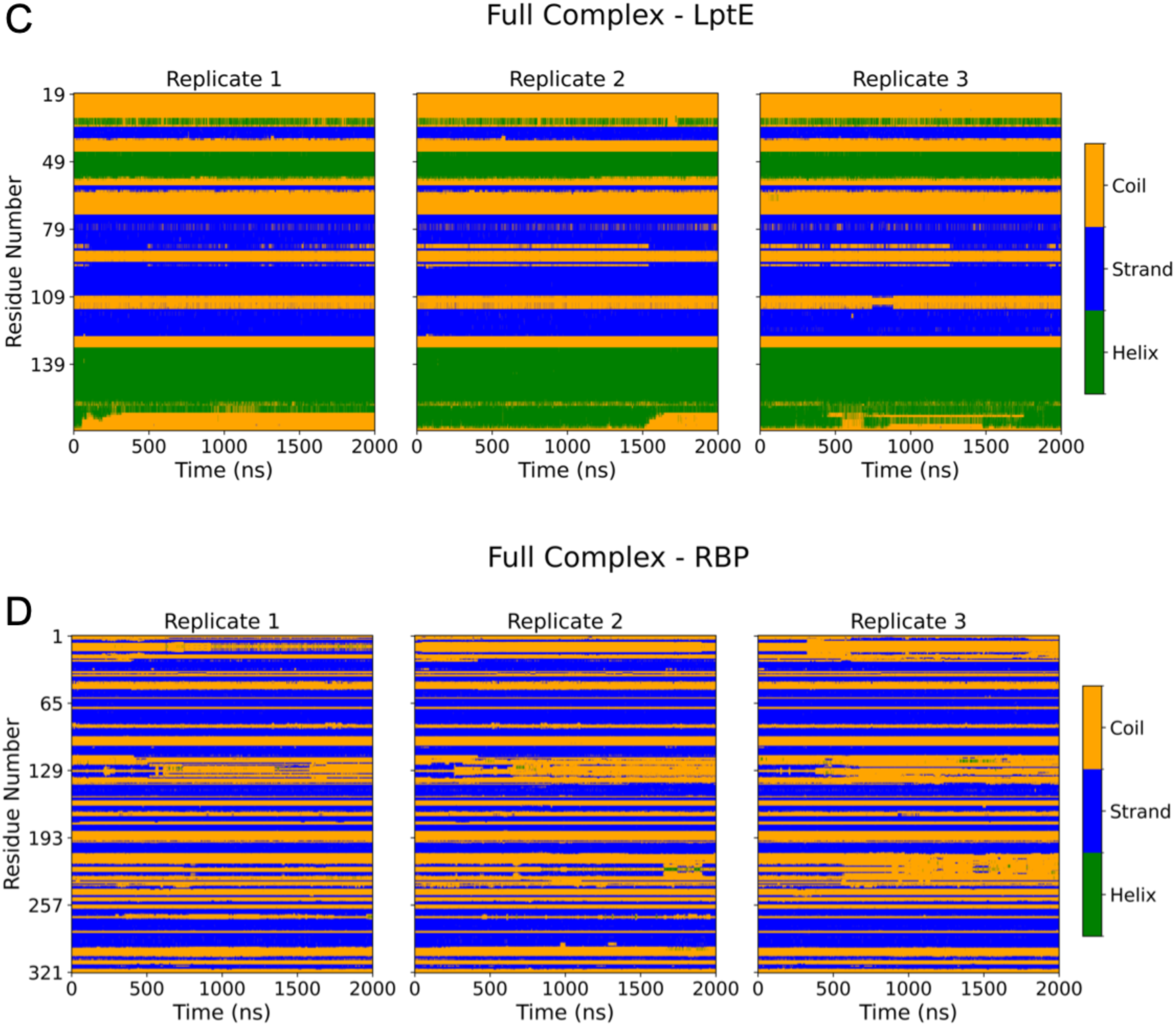
Stable secondary structure in the SfLptDE-RBP_Oeko_ complex. (A) Molecular dynamics simulations (3 x 2 μs) showing the overall secondary structure for each of the three proteins, as analysed by DSSP. (B-D) Sequence-specific secondary structure for LptD (B), LptE (C) and RBP_Oeko_ (D).

**Fig. S7.**
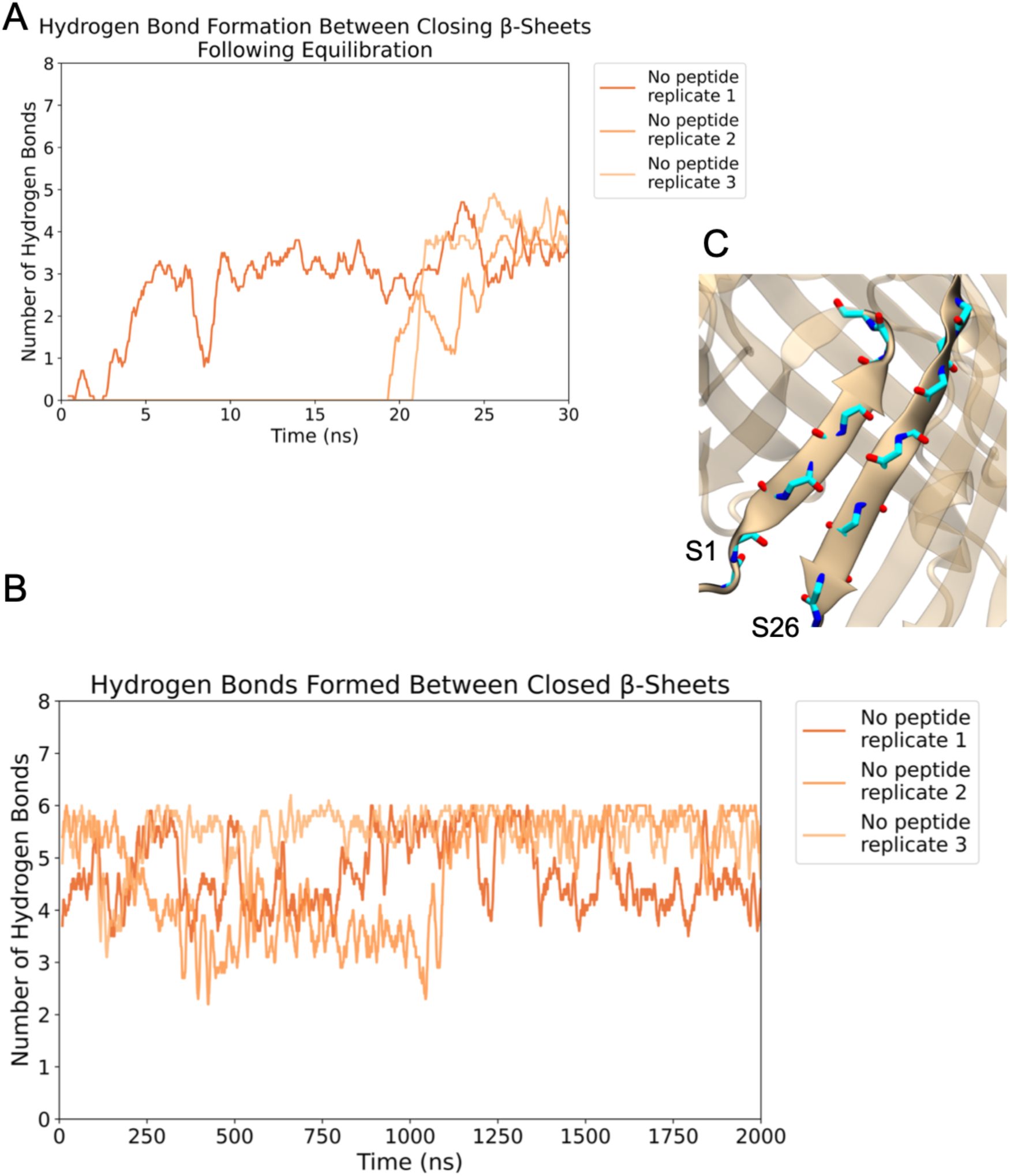
*In silico* removal of the S0 peptide results in fast and stable lateral gate closure. (A,B) Initial 30 ns and complete trajectories of the three simulations of SfLptDE-RBP_Oeko_ following S0 removal. The traces in (B) are rolling averages over 10 ns. (C) Representative snapshot of the closed lateral gate after S0 removal. Residue side chains are not shown for clarity.

**Fig. S8.**
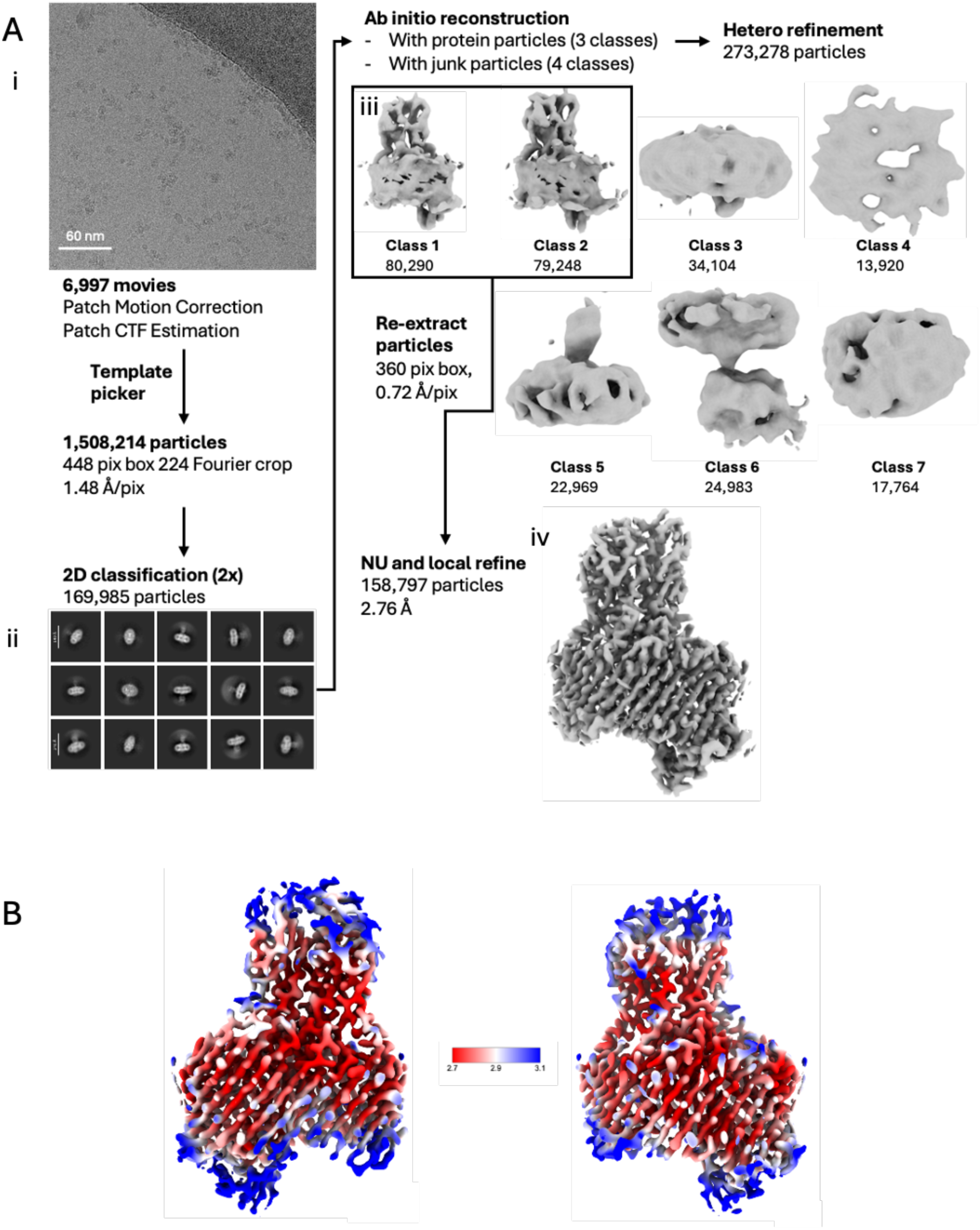
Cryo-EM data processing for the *Sf*LptD-RBP dataset 2. (A) Workflow summary. 6,997 movies were collected and imported into CryoSPARC. After motion correction and CTF estimation, low-quality micrographs were excluded. Particles were extracted with Fourier cropping and subjected to two rounds of 2D classification (i, example micrograph; ii, representative 2D classes) followed by ab-initio reconstruction and heterogenous refinement using protein and junk volumes (iii, volumes). particles were re-extracted at full resolution for non-uniform and local refinement, yielding a final 2.76Å map (iv, final volume). (B) Local resolution estimation of final map.

**Fig. S9.**
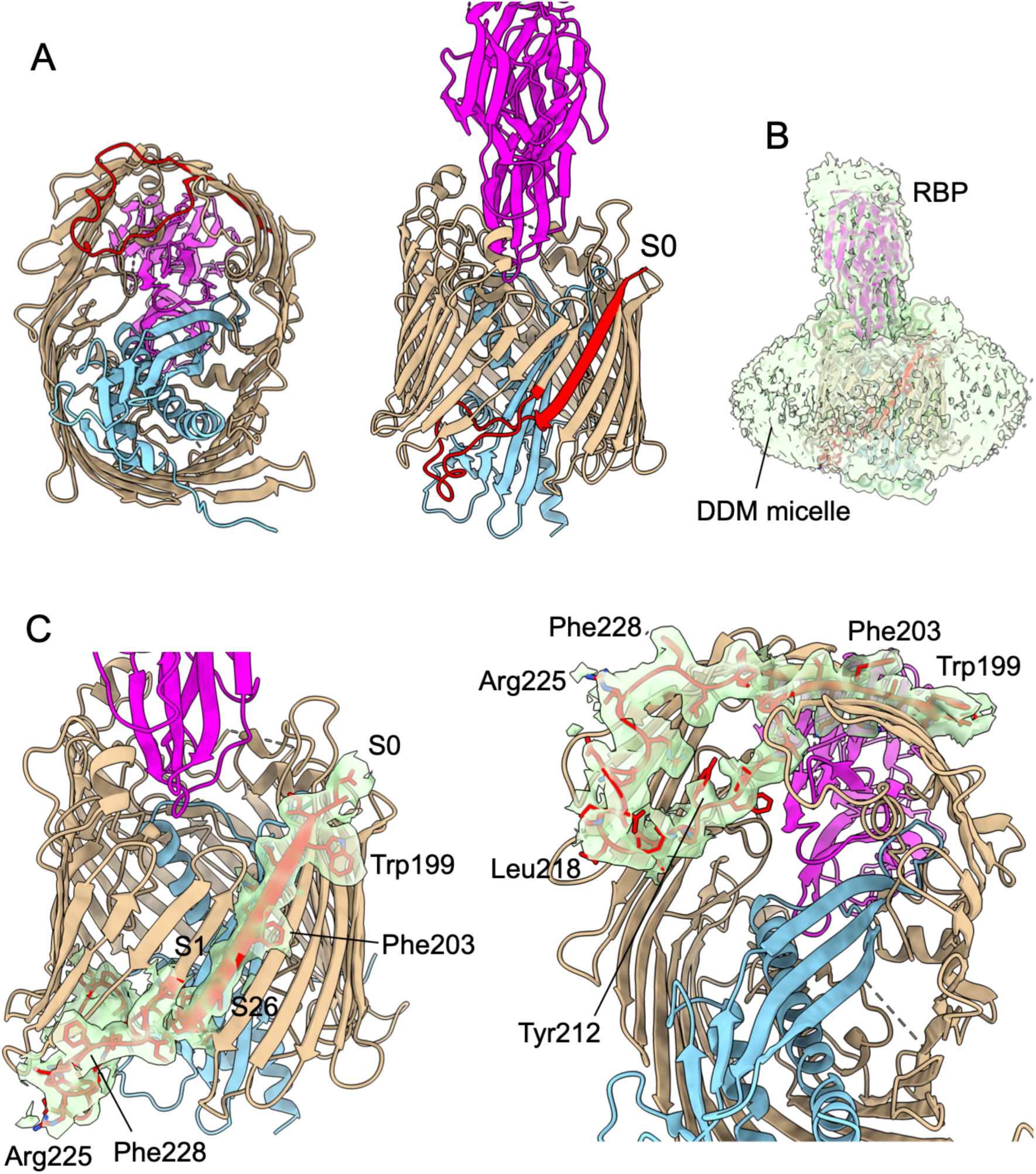
The lateral gate β-strand S0 is part of the jellyroll domain. (A) Cartoons from the periplasmic space (left) and from the front, showing LptD segment Glu197-Asn232 in red. (B) Low contour map of dataset 2. (C) Cartoons including carved low contour maps (as in B) for the Glu197-Asn232 segment of LptD. See also Movie S2.

**Fig. S10.**
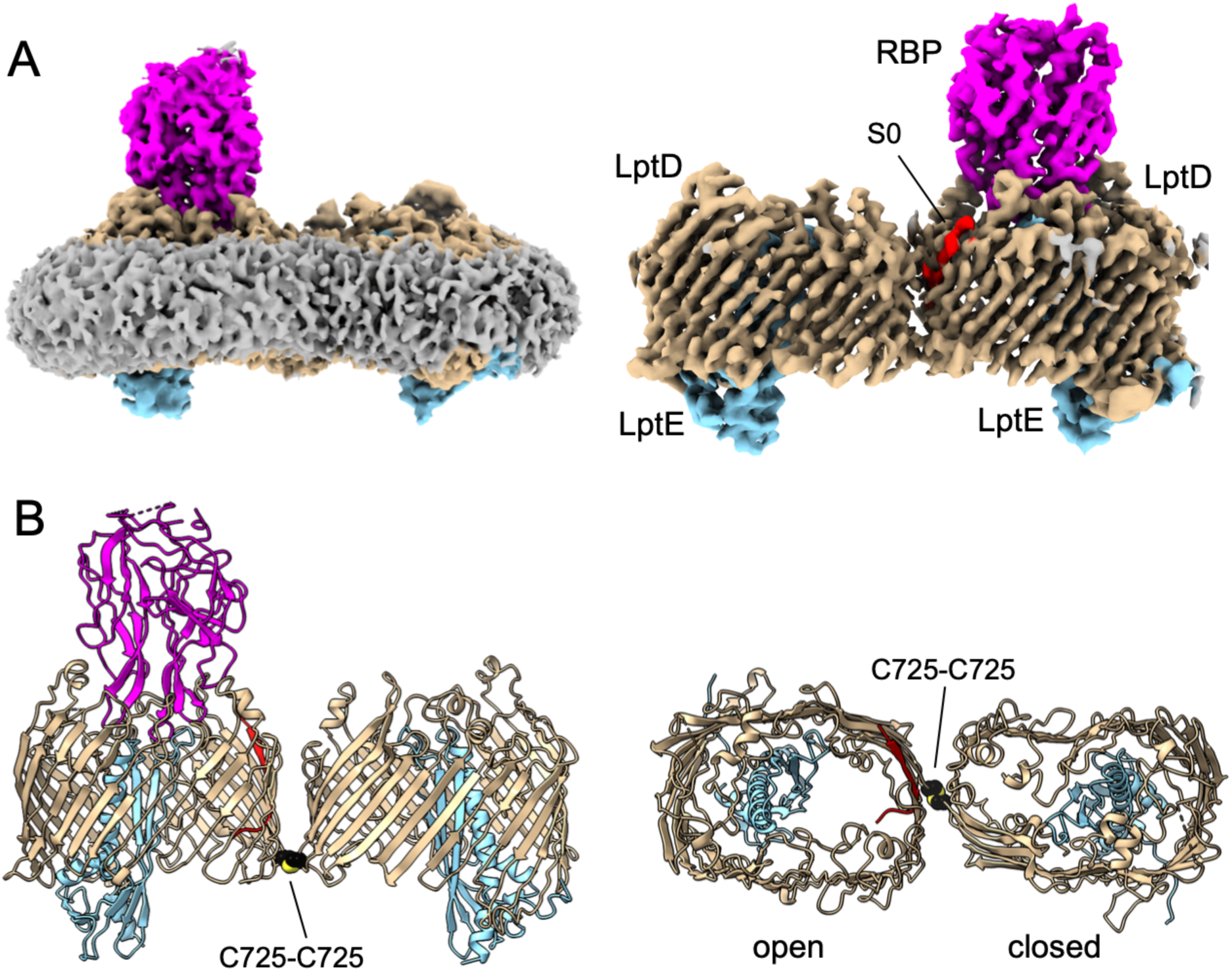
Structure of dimeric LptDE. (A) Low (left) and high contour maps of LptDE (dataset 1). (B) Cartoons viewed from the OM plane (left) and from the extracellular side. The intermolecular Cys725-Cys725 disulphide bond between LptD protomers is shown as space-filling models (carbons, black; sulphurs yellow). RBP is not shown in the extracellular view for clarity.

**Fig. S11.**
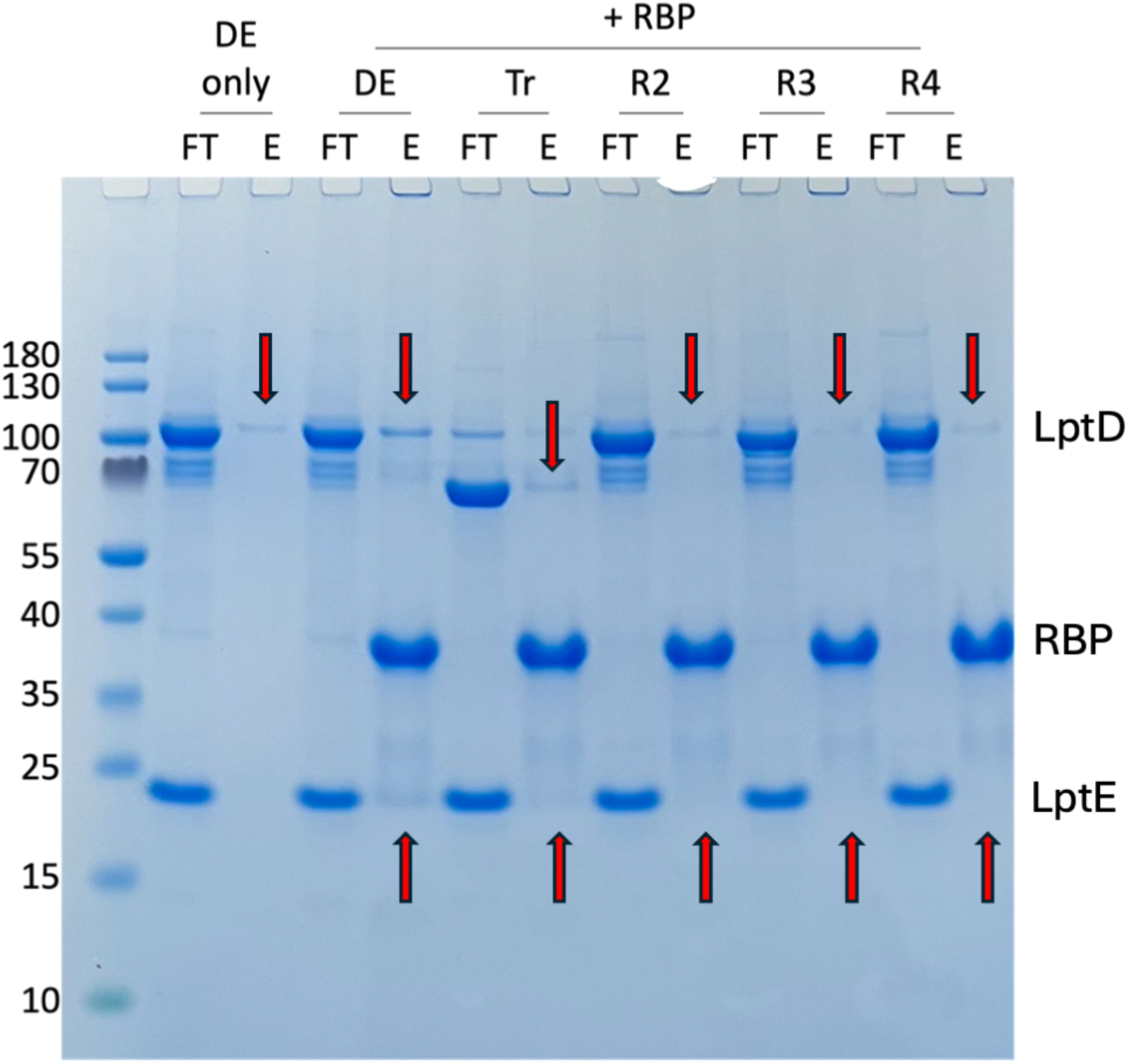
Pulldown experiments for LptDE +/− RBP incubations. Non-tagged WT and variant LptDE complexes were incubated with 3-fold molar excess of His-tagged RBP for 48 h at 4 C° and subjected to batch IMAC. Only flowthrough (FT) and elution (E) fractions are shown. The arrows indicate the expected positions of LptD and LptE within the elution fractions. DE, wild type LptDE; Tr, truncated LptD without the jellyroll domain (τι26-201); R2, LptD Y671N; R3, LptD D352Y; R4, LptD Thr385-Gln393 x 2 (duplication).

**Fig. S12.**
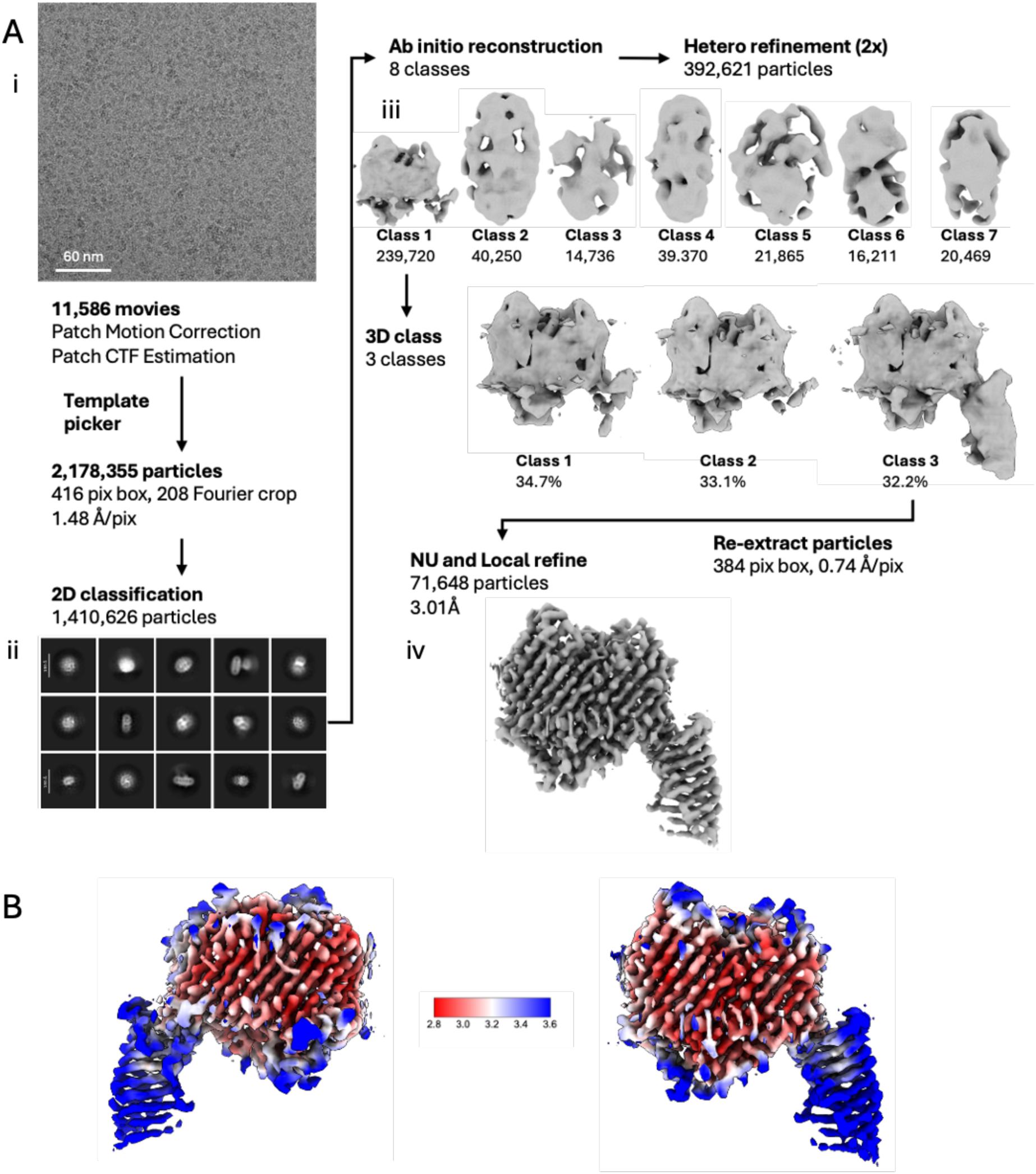
Cryo-EM data processing for *Sf*LptED-*Ec*LptM complex. (A) Workflow summary. 11,586 movies were collected and imported into CryoSPARC. After motion correction and CTF estimation, low-quality micrographs were excluded. Particles were extracted with Fourier cropping and subjected to two rounds of 2D classification (i, example micrograph; ii, representative 2D classes) followed by ab-initio reconstruction and heterogenous refinement using protein and junk volumes (iii, volumes). Particles were re-extracted at full resolution for non-uniform and local refinement, yielding a final 3.01Å map (iv, final volume). (B) Local resolution estimation of final map.

**Fig. S13.**
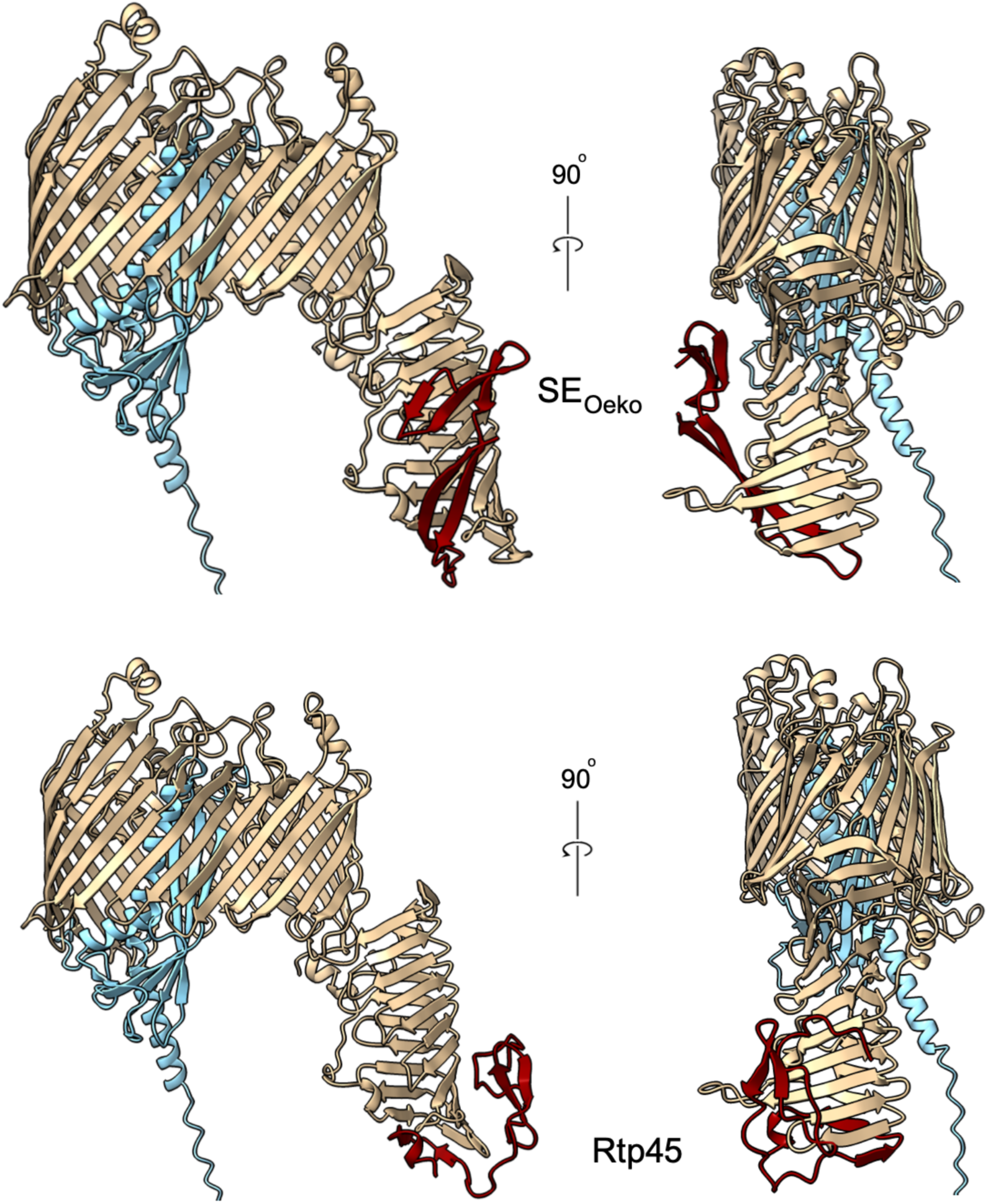
AF3 models of complexes between LptDE and SE_Oeko_ (top) and Rtp45 (bottom), viewed in the OM plane. LptD is coloured tan, LptE light blue and the SE proteins maroon.

**Table S1.** Cryo-EM data collection and processing parameters.

|  | **DE-RBP collection 1** |  | **DEM-RTP45** | **DEM** | **DE-RBP collection 2** |
| --- | --- | --- | --- | --- | --- |
| **Data collection** |  |  |  |  |  |
| Microscope | Titan Krios |  | Titan Krios | Titan Krios | Titan Krios |
| Detector | Falcon 4i (counting) |  | Falcon 4i (counting) | Falcon 4i (Selectris 10eV) | Falcon 4i (Selectris 10eV) |
| Voltage (kV) | 300 |  | 300 | 300 | 300 |
| Pixel size (Å) | 0.74 |  | 0.74 | 0.74 | 0.74 |
| Magnification | 165,000x |  | 165,000x | 165,000x | 165,000x |
| Total dose (e<sup>-</sup>/Å<sup>2</sup>) | 35.6 |  | 50.6 | 50 | 50 |
| Defocus range (µm) | -0.8 to -2.3 |  | -0.8 to -2.3 | -0.8 to -2.0 | -0.8 to -2.0 |
| Number of movies | 10,744 |  | 11,586 | 7,277 | 6,997 |
| **Image Processing** | **Class 1** | **Class 2** |  |  |  |
| Initial number of particles | 1,902,801 | 1,902,801 | 2,178,355 | 1,081,511 | 1,508,214 |
| Final number of particles | 134,671 | 55,494 | 71,648 | 112,032 | 158,797 |
| Map resolution (Å) | 2.81 | 3.20 | 3.01 | 2.78 | 2.76 |
| **Refinement** | **Monomer** | **Dimer** |  |  |  |
| MolProbity score | 1.01 | 2.5 | 1.47 | 1.6 | 1.72 |
| Clash score | 2.17 | 5.75 | 2.34 | 2.71 | 5.35 |
| Geometry |  |  |  |  |  |
| Bad bonds (%) | 0 | 0 | 0 | 0 | 0 |
| Bad angles (%) | 0 | 0.01 | 0 | 0 | 0 |
| Stereochemistry |  |  |  |  |  |
| Ramachandran favoured (%) | 98.01 | 87.79 | 94.67 | 95.55 | 95.38 |
| Ramachandran outliers (%) | 0 | 0.12 | 0.11 | 0 | 0 |
| Favoured rotamers (%) | 91.9 | 84.16 | 91.87 | 89.69 | 71.14 |
| Poor rotamers (%) | 1.07 | 5.85 | 1.33 | 2.01 | 1.4 |
| PDB | 9RPT | 9RPW | 9RPS | 9RPR | 9RQI |
| EMDB | EMD-54171 | EMD-54173 | EMD-54170 | EMD-54169 | EMD-54175 |

**Table S2.** MIC values for the L– and D-variant of the S0 peptide against a panel of wild-type and weakened *E. coli* strains with attenuated outer membranes. The panel of strains were chosen to include knockouts of non-essential genes involved in OMP biogenesis (Δ*surA* is a chaperone responsible for correct folding of the BAM complex), attenuated LPS production (Δ*waaD*) (3) and correct folding and disulfide bond formation of LptDE (Δ*dsbA*, Δ*dsbC*, Δ*lptM*) (4) GKCW102 is a hyperporinated strain with an inducible pore (BW25113 attTn7::mini-Tn7T Kmr araC ParaBAD fhuAΔC/Δ4L) that increases OM permemability, and GKCW101 is the control for this (BW25113 attTn7::mini-Tn7T Kmr araC ParaBAD MCS) (5) DC2 is an antibiotic-hypersusceptible E. coli mutant (6) NT = not tested, number of experimental repeats are shown in brackets.

| Bacterial Strain | MHB |  |
| --- | --- | --- |
|  | S0 L-form | S0 D-form |
| **E. coli Imp4213** | 2 – 32 (14) | 16 – 64 (7) |
| **E. coli $\Delta SurA$** | >64 (3) | >64 (3) |
| **E. coli $\Delta WaaD$** | >64 (4) | >64 (1) |
| **E. coli GKCW101** | >64 (4) | >64 (4) |
| **E. coli GKCW102** | >64 (4) | 64 - >64 (3) |
| **E. coli $\Delta DsbA$** | >64 (1) | >64 (1) |
| **E. coli $\Delta DsbC$** | >64 (1) | >64 (1) |
| **E. coli $\Delta LptM$** | >64 (1) | >64 (1) |
| **E. coli DC2** | >64 (1) | >64 (1) |
| **E. coli ATCC25922** | >64 (1) | >64 (1) |

**Movie S1.** The LptDEM structure. Map and cartoon showing bound LptM (green) within the LptD lumen.

**Movie S2.** LptDE with bound RBP_Oeko_, emphasising the connection between LptD strands S0 and S1 (red). Density from dataset 2 is shown for the connecting segment.

**Movie S3.** LptDE with bound RBP_Oeko_, emphasising the difference between the closed and open states of LptD. The movie starts with a density map (dataset 2) followed by a cartoon. Following the top view, RBP (magenta) is removed, and the cartoon is morphed into the closed state, with the S0 segment (red) occupying its position within the jellyroll domain. A surface view of the closed state follows. Subsequently, the closed state morphs back into the open state and ends with a surface view (RBP not shown for clarity).

**Movie S4.** Rtp45 binding to LptDEM. Movie starts with a cartoon of LptDE from the LptDEM dataset (LptD in tan, LptE in blue). The barrel is clipped open and morphed into the LptDE conformation from the LptDEM-Rtp45 complex and Rtp45 (maroon) moves into position. A surface representation of RBP (magenta), from the SfLptDE-RBP_Oeko_ complex is displayed to illustrate clashes with EL4 (highlighted in green). The map from the LptDEM-Rtp45 dataset is then shown. LptM is hidden for clarity.

## References

1. C. Anastassopoulou, S. Ferous, A. Petsimeri, G. Gioula, A. Tsakris, Phage-Based Therapy in Combination with Antibiotics: A Promising Alternative against Multidrug-Resistant Gram-Negative Pathogens. Pathogens 13, 896 (2024).

2. J. Bertozzi Silva, Z. Storms, D. Sauvageau, Host receptors for bacteriophage adsorption. FEMS Microbiol Lett 363, fnw002 (2016).

3. B. van den Berg, et al., Structural basis for host recognition and superinfection exclusion by bacteriophage T5. Proceedings of the National Academy of Sciences 119 (2022).

4. S. Degroux, G. Effantin, R. Linares, G. Schoehn, C. Breyton, Deciphering Bacteriophage T5 Host Recognition Mechanism and Infection Trigger. J Virol 97 (2023).

5. R. Linares, et al., Structural basis of bacteriophage T5 infection trigger and *E. coli* cell wall perforation. Sci Adv 9 (2023).

6. A. Silale, B. van den Berg, TonB-Dependent Transport Across the Bacterial Outer Membrane. Annu Rev Microbiol 77, 67–88 (2023).

7. R. Manley, et al., Resistance to bacteriophage incurs a cost to virulence in drug-resistant Acinetobacter baumannii. J Med Microbiol 73 (2024).

8. K. E. Kortright, R. E. Done, B. K. Chan, V. Souza, P. E. Turner, Selection for Phage Resistance Reduces Virulence of Shigella flexneri. Appl Environ Microbiol 88, e0151421 (2022).

9. J. Gurney, S. P. Brown, O. Kaltz, M. E. Hochberg, Steering Phages to Combat Bacterial Pathogens. Trends Microbiol 28, 85–94 (2020).

10. M. Braun, T. J. Silhavy, Imp/OstA is required for cell envelope biogenesis in Escherichia coli. Mol Microbiol 45, 1289–1302 (2002).

11. T. Wu, et al., Identification of a protein complex that assembles lipopolysaccharide in the outer membrane of Escherichia coli. Proc Natl Acad Sci U S A 103, 11754–11759 (2006).

12. S.-S. Chng, L. S. Gronenberg, D. Kahne, Proteins Required for Lipopolysaccharide Assembly in *Escherichia coli* Form a Transenvelope Complex. Biochemistry 49, 4565–4567 (2010).

13. S. Narita, H. Tokuda, Biochemical characterization of an ABC transporter LptBFGC complex required for the outer membrane sorting of lipopolysaccharides. FEBS Lett 583, 2160–2164 (2009).

14. P. Sperandeo, et al., New Insights into the Lpt Machinery for Lipopolysaccharide Transport to the Cell Surface: LptA-LptC Interaction and LptA Stability as Sensors of a Properly Assembled Transenvelope Complex. J Bacteriol 193, 1042–1053 (2011).

15. L. Törk, C. B. Moffatt, T. G. Bernhardt, E. C. Garner, D. Kahne, Single-molecule dynamics show a transient lipopolysaccharide transport bridge. Nature 623, 814–819 (2023).

16. S. Okuda, D. J. Sherman, T. J. Silhavy, N. Ruiz, D. Kahne, Lipopolysaccharide transport and assembly at the outer membrane: the PEZ model. Nat Rev Microbiol 14, 337–345 (2016).

17. S. Qiao, Q. Luo, Y. Zhao, X. C. Zhang, Y. Huang, Structural basis for lipopolysaccharide insertion in the bacterial outer membrane. Nature 511, 108–111 (2014).

18. X. Li, Y. Gu, H. Dong, W. Wang, C. Dong, Trapped lipopolysaccharide and LptD intermediates reveal lipopolysaccharide translocation steps across the Escherichia coli outer membrane. Sci Rep 5, 11883 (2015).

19. I. Botos, et al., Structural and Functional Characterization of the LPS Transporter LptDE from Gram-Negative Pathogens. Structure 24, 965–976 (2016).

20. H. Dong, et al., Structural basis for outer membrane lipopolysaccharide insertion. Nature 511, 52–56 (2014).

21. E. Maffei, et al., Systematic exploration of Escherichia coli phage–host interactions with the BASEL phage collection. PLoS Biol 19, e3001424 (2021).

22. A. Wietzorrek, H. Schwarz, C. Herrmann, V. Braun, The Genome of the Novel Phage Rtp, with a Rosette-Like Tail Tip, IsHomologous to the Genome of Phage T1. J Bacteriol 188, 1419–1436 (2006).

23. B. A. Sampson, R. Misra, S. A. Benson, Identification and characterization of a new gene of Escherichia coli K-12 involved in outer membrane permeability. Genetics 122, 491–501 (1989).

24. M. Botte, et al., Cryo-EM structures of a LptDE transporter in complex with Pro-macrobodies offer insight into lipopolysaccharide translocation. Nat Commun 13, 1826 (2022).

25. Y. Yang, et al., LptM promotes oxidative maturation of the lipopolysaccharide translocon by substrate binding mimicry. Nat Commun 14, 6368 (2023).

26. Q. Luo, et al., Surface lipoprotein sorting by crosstalk between Lpt and Lol pathways in gram-negative bacteria. Nat Commun 16, 4357 (2025).

27. Y. Gu, et al., Lipopolysaccharide is inserted into the outer membrane through an intramembrane hole, a lumen gate, and the lateral opening of LptD. Structure 23, 496– 504 (2015).

28. G. Malojčić, et al., LptE binds to and alters the physical state of LPS to catalyze its assembly at the cell surface. Proceedings of the National Academy of Sciences 111, 9467–9472 (2014).

29. K. A. Black, et al., Resolution of a T1-Like Bacteriophage Outbreak by Receptor Engineering. Mol Biotechnol (2025). 10.1007/s12033-025-01453-1.

30. E. Krissinel, K. Henrick, Inference of Macromolecular Assemblies from Crystalline State. J Mol Biol 372, 774–797 (2007).

31. O. S. Smart, J. G. Neduvelil, X. Wang, B. A. Wallace, M. S. P. Sansom, HOLE: A program for the analysis of the pore dimensions of ion channel structural models. J Mol Graph 14, 354–360 (1996).

32. H. Zhang, et al., Genomic and biological insights of bacteriophages JNUWH1 and JNUWD in the arms race against bacterial resistance. Front Microbiol 15 (2024).

33. K. M. Kuo, J. Liu, A. Pavlova, J. C. Gumbart, Drug Binding to BamA Targets Its Lateral Gate. J Phys Chem B 127, 7509–7517 (2023).

34. D. Sun, et al., The discovery and structural basis of two distinct state-dependent inhibitors of BamA. Nat Commun 15, 8718 (2024).

35. N. Srinivas, et al., Peptidomimetic antibiotics target outer-membrane biogenesis in Pseudomonas aeruginosa. Science (1979) 327, 1010–1013 (2010).

36. J. A. Robinson, Folded Synthetic Peptides and Other Molecules Targeting Outer Membrane Protein Complexes in Gram-Negative Bacteria. Front Chem 7 (2019).

37. A. Schmidt, et al., The quantitative and condition-dependent Escherichia coli proteome. Nat Biotechnol 34, 104–110 (2016).

38. H. He, et al., A *Borrelia burgdorferi* LptD homolog is required for flipping of surface lipoproteins through the spirochetal outer membrane. Mol Microbiol 119, 752–767 (2023).

39. C. E. Cowles, Y. Li, M. F. Semmelhack, I. M. Cristea, T. J. Silhavy, The free and bound forms of Lpp occupy distinct subcellular locations in *Escherichia coli*. Mol Microbiol 79, 1168–1181 (2011).

40. A. Punjani, J. L. Rubinstein, D. J. Fleet, M. A. Brubaker, cryoSPARC: algorithms for rapid unsupervised cryo-EM structure determination. Nat Methods 14, 290–296 (2017).

41. A. Punjani, H. Zhang, D. J. Fleet, Non-uniform refinement: adaptive regularization improves single-particle cryo-EM reconstruction. Nat Methods 17, 1214–1221 (2020).

42. J. Jumper, et al., Highly accurate protein structure prediction with AlphaFold. Nature 596, 583–589 (2021).

43. E. F. Pettersen, et al., UCSF Chimera—A visualization system for exploratory research and analysis. J Comput Chem 25, 1605–1612 (2004).

44. P. D. Adams, et al., *PHENIX*: a comprehensive Python-based system for macromolecular structure solution. Acta Crystallogr D Biol Crystallogr 66, 213–221 (2010).

45. P. Emsley, B. Lohkamp, W. G. Scott, K. Cowtan, Features and development of Coot. Acta Crystallogr D Biol Crystallogr 66, 486–501 (2010).

46. Schrödinger LLC, “The PyMOL Molecular Graphics System, Version 2.5 Schrödinger, LLC.” (2015).

47. S. Jo, T. Kim, V. G. Iyer, W. Im, CHARMM-GUI: A web-based graphical user interface for CHARMM. J Comput Chem 29, 1859–1865 (2008).

48. E. L. Wu, et al., CHARMM-GUI membrane builder toward realistic biological membrane simulations. J Comput Chem 35, 1997–2004 (2014).

49. J. Lee, et al., CHARMM-GUI Membrane Builder for Complex Biological Membrane Simulations with Glycolipids and Lipoglycans. J Chem Theory Comput 15, 775–786 (2019).

50. G. Bussi, D. Donadio, M. Parrinello, Canonical sampling through velocity rescaling. Journal of Chemical Physics 126, 14101 (2007).

51. M. Bernetti, G. Bussi, Pressure control using stochastic cell rescaling. Journal of Chemical Physics 153, 114107 (2020).

52. B. Hess, P-LINCS: A parallel linear constraint solver for molecular simulation. J Chem Theory Comput 4, 116–122 (2008).

53. U. Essmann, et al., A smooth particle mesh Ewald method. J Chem Phys 103, 8577– 8593 (1995).

54. J. Huang, et al., CHARMM36m: an improved force field for folded and intrinsically disordered proteins. Nature Methods 2016 14:1 14, 71–73 (2016).

55. M. J. Abraham, et al., GROMACS: High performance molecular simulations through multi-level parallelism from laptops to supercomputers. SoftwareX 1–2, 19–25 (2015).

56. O. S. Smart, J. M. Goodfellow, B. A. Wallace, The pore dimensions of gramicidin A. Biophys J 65, 2455 (1993).

57. N. Michaud-Agrawal, E. J. Denning, T. B. Woolf, O. Beckstein, MDAnalysis: A toolkit for the analysis of molecular dynamics simulations. J Comput Chem 32, 2319– 2327 (2011).

58. R. J. Gowers, et al., MDAnalysis: A Python Package for the Rapid Analysis of Molecular Dynamics Simulations. scipy 98–105 (2016). 10.25080/MAJORA-629E541A-00E.

59. W. Kabsch, C. Sander, Dictionary of protein secondary structure: Pattern recognition of hydrogen-bonded and geometrical features. Biopolymers 22, 2577–2637 (1983).

60. R. T. McGibbon, et al., MDTraj: A Modern Open Library for the Analysis of Molecular Dynamics Trajectories. Biophys J 109, 1528–1532 (2015).

61. E. N. Baker, R. E. Hubbard, Hydrogen bonding in globular proteins. Prog Biophys Mol Biol 44, 97–179 (1984).

## Supplementary references

1. A. Punjani, J. L. Rubinstein, D. J. Fleet, M. A. Brubaker, cryoSPARC: algorithms for rapid unsupervised cryo-EM structure determination. Nat Methods 14, 290–296 (2017).

2. A. Punjani, H. Zhang, D. J. Fleet, Non-uniform refinement: adaptive regularization improves single-particle cryo-EM reconstruction. Nat Methods 17, 1214–1221 (2020).

3. T. Baba, et al., Construction of *Escherichia coli* K-12 in-frame, single-gene knockout mutants: the Keio collection. Mol Syst Biol 2 (2006).

4. Y. Yang, et al., LptM promotes oxidative maturation of the lipopolysaccharide translocon by substrate binding mimicry. Nat Commun 14, 6368 (2023).

5. G. Krishnamoorthy, et al., Breaking the Permeability Barrier of Escherichia coli by Controlled Hyperporination of the Outer Membrane. Antimicrob Agents Chemother 60, 7372–7381 (2016).

6. D. Clark, Novel antibiotic hypersensitive mutants of *Escherichia coli* genetic mapping and chemical characterization. FEMS Microbiol Lett 21, 189–195 (1984).

